# Developmental regulators FlbE/D orchestrate the polarity site-to-nucleus dynamics of the fungal bZIP FlbB

**DOI:** 10.1101/467563

**Authors:** Ainara Otamendi, Elixabet Perez-de-Nanclares-Arregi, Elixabet Oiartzabal, Marc S. Cortese, Eduardo A. Espeso, Oier Etxebeste

## Abstract

Permanently polarized cells have developed transduction mechanisms linking polarity-sites with gene regulation in the nucleus. In neurons, one mechanism is based on long-distance retrograde migration of transcription factors (TFs). *Aspergillus nidulans* FlbB is the only known fungal TF shown to migrate retrogradely to nuclei from the polarized region of fungal cells known as hyphae. There, FlbB controls developmental transitions by triggering the production of asexual multicellular structures. FlbB dynamics in hyphae is orchestrated by regulators FlbE and FlbD. At least three FlbE domains are involved in the acropetal transport of FlbB, with a final MyoE/actin filament-dependent step from the subapex to the apex. Experiments employing a T2A viral peptide-containing chimera (FlbE::mRFP::T2A::FlbB::GFP) suggest that apical FlbB/FlbE interaction is inhibited in order to initiate a dynein-dependent FlbB transport to nuclei. FlbD controls the nuclear accumulation of FlbB through a cMyb domain and a C-terminal LxxLL motif. Overall, results elucidate a highly dynamic pattern of FlbB interactions, which enable timely developmental induction. Furthermore, this system establishes a reference for TF-based long-distance signaling in permanently polarized cells.

## Introduction

The ability to adapt to changes in environmental conditions is a key feature of living organisms. With this aim, eukaryotic cells monitor the environment, receive signals from it, interiorize, amplify and integrate these cues, and finally convey the corresponding information to nuclei in the form of proteins (transcription factors; TFs) that are able of modifying gene expression patterns and, consequently, induce the adaptive cellular response. All these steps are carried out following a variety of molecular mechanisms that are generally classified as signal transduction pathways [1].

Permanent polarization of specific cell types such as neurons, pollen tubes or hyphae (see next paragraph) significantly increases the distance between polarity sites (i.e. growth cones in neurons or tips of hyphae) and nuclei, complicating the transduction of signals along this path [2–5]. Thus, permanently polarized cells have necessarily developed transduction mechanisms that are capable of covering the corresponding distances with speed and reliability. In neurons, these mechanisms are based mainly on calcium waves, but also on the retrograde transport of macromolecular complexes [5–7]. Retrograde transport of macromolecular complexes are used to control key neuronal processes such as the response to injury [8]. The main messengers in those complexes can be kinases which ultimately transfer the signal to a TF in the nuclear periphery or inside the nucleus, or TFs themselves that are able to migrate basipetally from the polarity site to the nucleus and directly modify gene expression.

Hyphae are the characteristic cell type of filamentous fungi. These permanently-polarized structures elongate by pulsed extension of the tip apex [9] that is dependent on receiving plasma membrane and cell-wall materials that are transported first on microtubules (MT) and then on actin filaments [10,11]. The fast polar growth of fungal hyphae increases turgor pressure impinged on the substrate, facilitating its efficient colonization. Hyphae also sense the environment and vary their growth direction in response to different stimuli such as chemical, topographical or electrical signals [12–17]. Under unfavorable growth conditions [18–20] and/or in response to specific chemical signals [21], developmental transitions are triggered in hyphae, producing sexual or asexual spores depending on the stimulus [22]. Asexual spores are mitotic spores constituting the prevalent mechanism for dissemination of fungi. Recent findings have shown the existence of signaling complexes retrogradely transiting from the tip of hyphae to nuclei [3,5,23–26]. Although a limited number of them have been characterized, these mechanisms are based, as in neurons, either on kinase modules or the direct basipetal transport of TFs. These factors control stress responses as well as the sexual and asexual multicellular developmental cycles of filamentous fungal species such as *Aspergillus nidulans*.

This ascomycete is the preferential reference organism used in the study of the genetic and molecular control of fungal asexual development [27]. Most of the developmental transitions leading this fungus to the production of asexual spores, known as conidia, are controlled by the central developmental pathway (CDP) [20]. *brlA* is the first CDP gene and, thus, many signal transduction pathways activating or inhibiting conidiation converge at its promoter region so as to coordinately control its expression [28]. The upstream developmental activation (UDA) pathway is one of the main signal transduction pathways controlling *brlA* expression [29]. Three UDA TFs, FlbB, FlbC and FlbD, bind the *brlA* promoter [30,31] and control conidiation jointly with TFs from other pathways [28]. FlbC has been located in a sub-pathway parallel to that defined by FlbB and FlbD [31]. The regulatory activity of FlbB and FlbD is interdependent, since the former controls the expression of *flbD* but it cannot bind the promoter of *brlA* in the absence of FlbD [30].

The regulatory activity of FlbB strongly depends on its subcellular dynamics. FlbB is the first known fungal TF showing an apical localization [32]. Indeed, previous work showed that the FlbE-dependent tip localization of FlbB is a pre-requisite for timely control of *brlA* expression and that this TF is transported basipetally from the growth region to nuclei (hyphae of *A. nidulans* are multinucleate) [23]. In this work, we have delimited in space and time the role of UDA proteins FlbE and FlbD in regards to the control of FlbB dynamics. Through at least three domains, FlbE plays an essential role in the acropetal dynamics of FlbB towards the growing apex of the tip. Nevertheless, results suggest that FlbB/FlbE interaction is inhibited by an as yet unknown mechanism, initiating a tip-to-nucleus dynein- (and thus MT-) dependent basipetal migration of FlbB. FlbD positively controls the nuclear accumulation of FlbB through at least a highly conserved N-terminal cMyb transcriptional regulatory domain and a C-terminal LxxLL motif. Taking everything into consideration, results suggest that a precise sequence of interactions determines the directionality of FlbB dynamics, facilitating communication between the hyphal tip and nuclei, and consequently leading to timely coordination of the TFs that control the expresson of *brlA*.

## Results

### Constitutive upregulation of *flbE* increases the apical concentration of FlbB and induces conidiation in liquid culture

FlbB accumulates at the tip of vegetative hyphae and shows a concentration gradient in nuclei, with the highest concentration found in the most apical nucleus and steadily decreasing quantities in successive nuclei [32]. Constitutive expression of a GFP::FlbB chimera driven by the *gpdA*^*mini*^ promoter [33] increases the nuclear pool of FlbB, with all nuclei being filled with more or less equal amounts of the TF [23]. However, this excess in the nuclear pool of FlbB does not correlate with an increase in conidia production because it corresponds to a transcriptionally inactive form of this TF. The absence of the FlbB-interactor protein FlbE precludes accumulation of FlbB at the tip, linking the activation of FlbB to its transport to and/or accumulation at the growth region as well as the presence of FlbE [23,34].

To determine whether the apical concentration of FlbB is directly dependent on the quantity of FlbE available, we constitutively expressed FlbE::mRFP or FlbE::Stag fusions, each driven by the *gpdA*^*mini*^ promoter (integrated at the *flbE locus*; see Figure EV1A), in a *gpdA*^*mini*^::GFP::FlbB strain. Both dual over-expression (OE) strains showed a statistically significant increase in the apical fluorescence intensity of GFP::FlbB compared to the parental *gpdA*^*mini*^::GFP::FlbB strain (Figure 1A). The green fluorescence intensity ratio between the tip and the most apical nucleus increased significantly from 1.11 ± 0.24 in the control to 1.78 ± 0.45 in the FlbE::Stag strain and 1.83 ± 0.53 in the FlbE::RFP strain (n = 15 hyphae for each strain; p = 6.3 x 10^-6^ and 5.5 x 10^-5^, respectively). These results strongly suggest that the concentration of FlbB at the hyphal apex is tightly linked to the concentration of FlbE.

**Figure 1:**
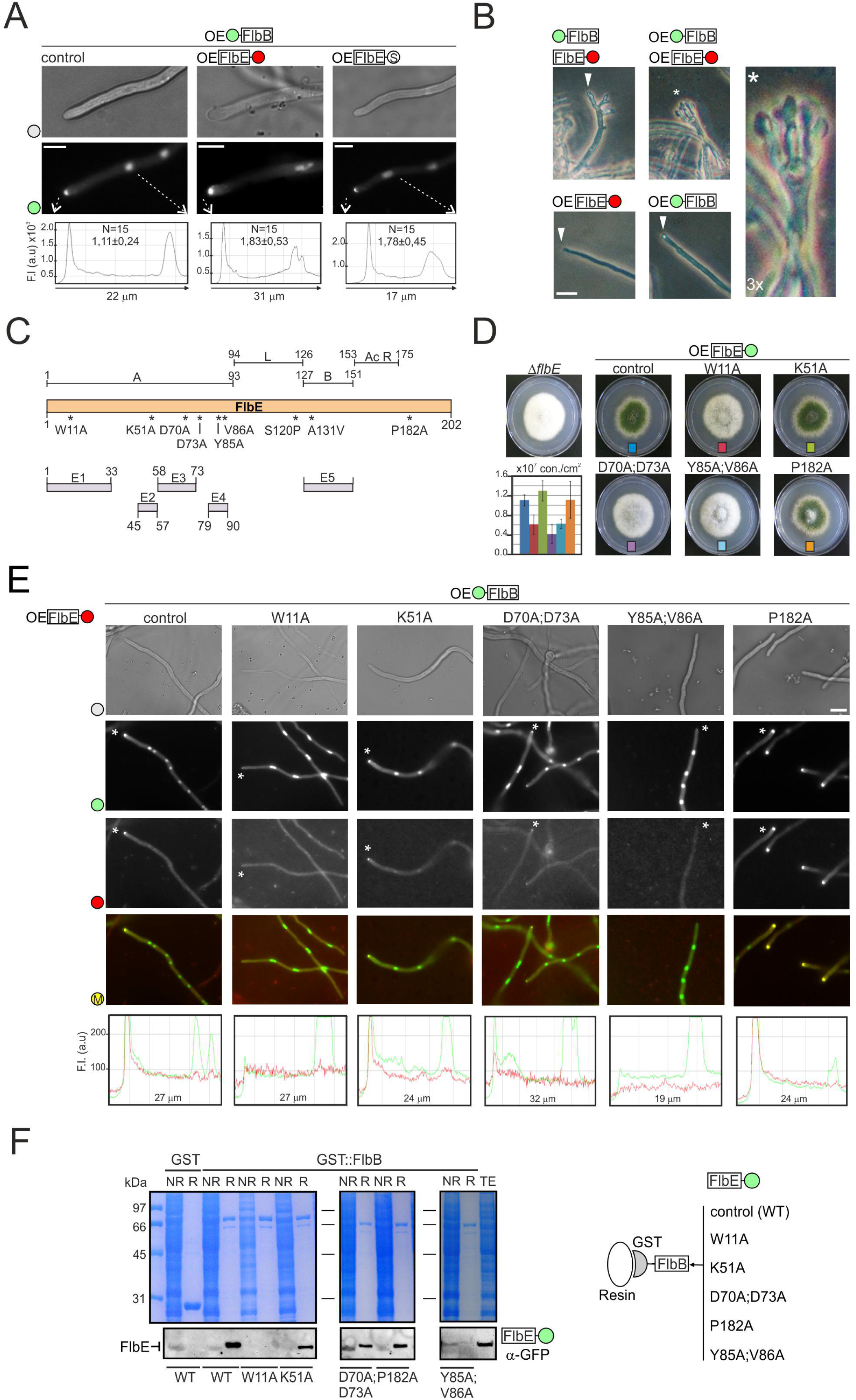
Role of FlbE domains in the apical localization of FlbB. A) Subcellular localization of GFP::FlbB in *gpdA*^*mini*^(OE)::GFP::FlbB (control), *gpdA*^*mini*^::GFP::FlbB;*gpdA*^*mini*^::FlbE::mRFP or *gpdA*^*mini*^::GFP::FlbB; *gpdA*^*mini*^::FlbE::Stag strains. The graphs below show the intensity of green fluorescence in the region delimited by the dotted arrows. The values show the fluorescence intensity ratio between the tip and the most apical nucleus, as the average of 15 measurements for each strain plus s.e.m. Scale bar = 5 µm B) Phenotype of 1) GFP::FlbB;FlbE::mCh, 2) *gpdA*^*mini*^::GFP::FlbB, 3) *gpdA*^*mini*^::FlbE::mRFP and 4) *gpdA*^*mini*^::GFP::FlbB;*gpdA*^*mini*^::FlbE::mRFP strains after 26 hours of culture in liquid AMM. The double-*gpdA*^*mini*^ strain developed conidiophores (see the asterisk), which can be seen in the 3x amplification on the right. Scale bar = 10 µm. C) Predicted functional domains within FlbE, based on the analyses described by [34] (up) and [35] (down). D) Phenotype of point *flbE\** mutants in AMM plates after 72 hours of culture at 37 °C. The graph shows conidia counts per cm^2^ for each mutant. Values are given as the mean of three replicates plus s.e.m. E) Subcellular localization of *gpdA*^*mini*^::GFP::FlbB (wild-type form) and *gpdA*^*mini*^::FlbE*::RFP (mutant forms) in vegetative hyphae (the wild-type form is shown as control). Scale bar = 5 µm. The graphs below show red and green fluorescence intensities (arbitrary units) in hyphal segments covering the region between the tip and the most apical nucleus (asterisks indicate which hyphae have been analyzed). M: merged. F) Co-immunoprecipitation assays using bacterially expressed GST (negative control) and GST::FlbB forms and crude extracts from strains expressing wild-type or mutant *gpdA*^*mini*^::FlbE::GFP forms W11A, K51A, D70A;D73A, Y85A;V86A and P182A. Polyacrilamide gels stained with Coomassie blue are shown as controls. See also Figure EV1.

Next, we checked if this higher apical accumulation of FlbB and FlbE was accompanied by higher conidia production and/or premature induction of conidiation (Figure 1B and Figure EV1B). On solid medium, wild-type strains conidiate because hyphae are exposed to the air environment [20]. The three strains produced similar amounts of conidia (n = 3 for each strain; p =0.41 and 0.59, when mRFP- or S-tagged strains were compared to the reference strain) (Figure EV1B). Clear differences arose, however, after 26 hours of culture in liquid medium compared to the following three reference strains (Figure 1B): 1) a strain expressing GFP::FlbB and FlbE::mCherry chimeras, each driven by its respective native promoter; 2) a strain expressing a *gpdA*^*mini*^-driven GFP::FlbB chimera; or 3) a strain expressing a *gpdA*^*mini*^-driven FlbE::RFP chimera. While, as expected, reference strains formed only vegetative hyphae (triangles in Figure 1B), the double-*gpdA*^*mini*^ strain produced conidiophores (the asexual structures bearing conidia) composed of all the characteristic cell types (asterisk in Figure 1B). Results strongly suggest that the apical accumulation of FlbB is directly dependent on FlbE concentration, and that a higher accumulation of the FlbB/FlbE signaling complex at the tip correlates, under certain growth conditions, with the ability to induce conidiation prematurely.

### Domain analysis of FlbE

Due to the key role of FlbE in the apical accumulation of FlbB and, consequently, the control of conidiation, we proceeded with a deeper characterization of functional domains in FlbE. Preliminary analysis of the FlbE sequence revealed the presence of two main domains (Figure 1C) [34]. Motif A spans from residue M1 to I93, and motif B resides between residues R127 and P151. Both regions were connected by a non-conserved linker sequence (94-126). The C-terminal region of FlbE, comprising residues from D153 to S202, showed a concentration of acidic, mainly aspartic, residues in the domain between D153 and D175, which are relatively conserved in most orthologs [34,35]. In contrast, the region between G176 and S202 showed no conservation among FlbE orthologs.

A more detailed HMM analysis divided motif A into four conserved regions,: E1 (1-33), E2 (45-57), E3 (58-73) and E4 (79-90) (domain B, positions 127-151, was renamed as E5 in this scheme) (Figure 1C) [35]. Specific residues within conserved domains were selected for alanine replacement. Due to its absolute conservation in FlbE orthologs [35], W11 was selected to study the role of domain E1. As a potential ubiquitination target [36], K51 was selected within domain E2. D70 and D73 were also mutated as they might contribute to a polyproline helix structure in domain E3. Y85 and V86, respectively, which are located within a predicted hydrophobic cluster in domain E4 and are highly conserved within FlbE orthologs, were also selected. Finally, and due to the characteristic conformational restrictions imposed by prolines, P182 was selected in the poorly-conserved C-terminal region. We had previously described that mutation S120P within the linker region and an A131V substitution within domain E5 both caused delocalization of FlbE from the tip and, consequently, a *fluffy* aconidial phenotype [34]. Thus, they were not included in the current analysis.

First, a strain expressing a *gpdA*^*mini*^-driven FlbE::GFP chimera was generated following the same procedure as that shown in Figure EV1A. Using its genomic DNA as a template, we proceeded with the site-directed mutagenesis approach described in Figure EV1C, in order to generate strains of *A. nidulans* expressing the *gpdA*^*mini*^-driven mutant FlbE::GFP forms described above. The replacement of the targeted amino acid and accuracy of *flbE* sequence were confirmed by sequencing of the complete open reading frame. The phenotype of the strains and the subcellular localization of the mutant chimeras were then analyzed (Figure 1D; Figure 1E for mRFP-tagged mutant FlbE forms; Figure EV1D for GFP-tagged counterparts). Mutations K51A and P182A did not alter conidia production (p > 0.05 compared to the parental wild-type strain; n = 3 replicates for each strain) nor FlbB/FlbE localization. The fact that P182A mutation did not inhibit conidia production or FlbE/FlbB localization probably reflects that hypothetic folding induced by this residue is not essential for FlbE functionality.

Mutations W11A, D70A;D73A and Y85A;V86A caused an aconidial phenotype, with significantly reduced conidia production compared to the parental strain (Figure 1D). The wild-type *gpdA*^*mini*^::FlbE::GFP strain produced 1.1 x 10^7^ ± 1.2 x 10^6^ conidia/cm^2^ while W11A, D70A;D73A and Y85A;V86A mutants produced 4.1-6.2 x 10^6^ ± 0.9-2.0 x 10^6^ conidia/cm^2^ (p = 0.020, 0.006 and 0.010, respectively, in the three replicate experiments, with n = 3 replicates for each strain). Additionally, we observed differences in FlbE localization among these mutants. While W11A and Y85A;V86A mutations caused the absence of FlbE from the hyphal tip and a dispersion along the cytoplasm, *gpdA*^*mini*^-driven FlbE^(D70A;D73A)^::GFP (or –mRFP tagged) still accumulated at the tip (Figure 1E and Figure EV1D). However, due to the *fluffy* phenotype of the strain, it can be inferred that this apical form of FlbE is not fully functional or is not accumulated at the tip above a hypothetic threshold concentration (Figure 1D). In general, FlbB localization correlated with that of FlbE, being delocalized from the tip in W11A and Y85A;V86A mutants but not completely in the D70A:D73A mutant (Figure 1E; note also in Figure EV1E the localization of a GFP::FlbB chimera driven by the native *flbB* promoter in a strain co-expressing a *gpdA*^*mini*^-driven FlbE^(D70A;D73A)^::mRFP chimera). All *gpdA*^*mini*^-driven FlbE::GFP chimeras were detected by immunodetection and showed the same mobility on Western blots (Figure EV1F).

Finally, all mutant forms of FlbE were tested in immunoprecipitation assays against a bacterially expressed GST::FlbB form (Figure 1F). The results correlated with the localization of FlbE/FlbB. Those mutations within FlbE inhibiting the apical localization of FlbE/FlbB also showed inhibition of the *in vitro* interaction with GST::FlbB while those not affecting apical localization exhibited the interaction.

### Domain E1 is essential but insufficient for the apical localization of FlbB and FlbE

Next, we focused on domain E1 because this region was predicted to be a signal peptide in the AspGD (www.aspgd.org) database based on Interpro [37] search results. As a preliminary approach to investigate its possible function, the wild-type FlbE sequence was tagged with GFP at the N-terminus and expressed driven by the native *flbE* promoter (see the tagging procedure in Figure EV2A). N-terminal GFP tagging of FlbE caused an aconidial phenotype that was the consequence of the delocalization of FlbE from the tip (Figure 2A and 2B). As stated before, this result contrasts with the wild-type functionality observed for a C-terminal fluorescent tagged FlbE constructs (Figure 1 and EV1). A similar aconidial phenotype and delocalization of FlbE from the tip were observed in cells of a strain expressing the truncated FlbE^(34-201)^::GFP form lacking the putative signal peptide (Figure 2A, 2B and EV2B). These results show the importance of domain E1 in the function of FlbE in the cell.

**Figure 2:**
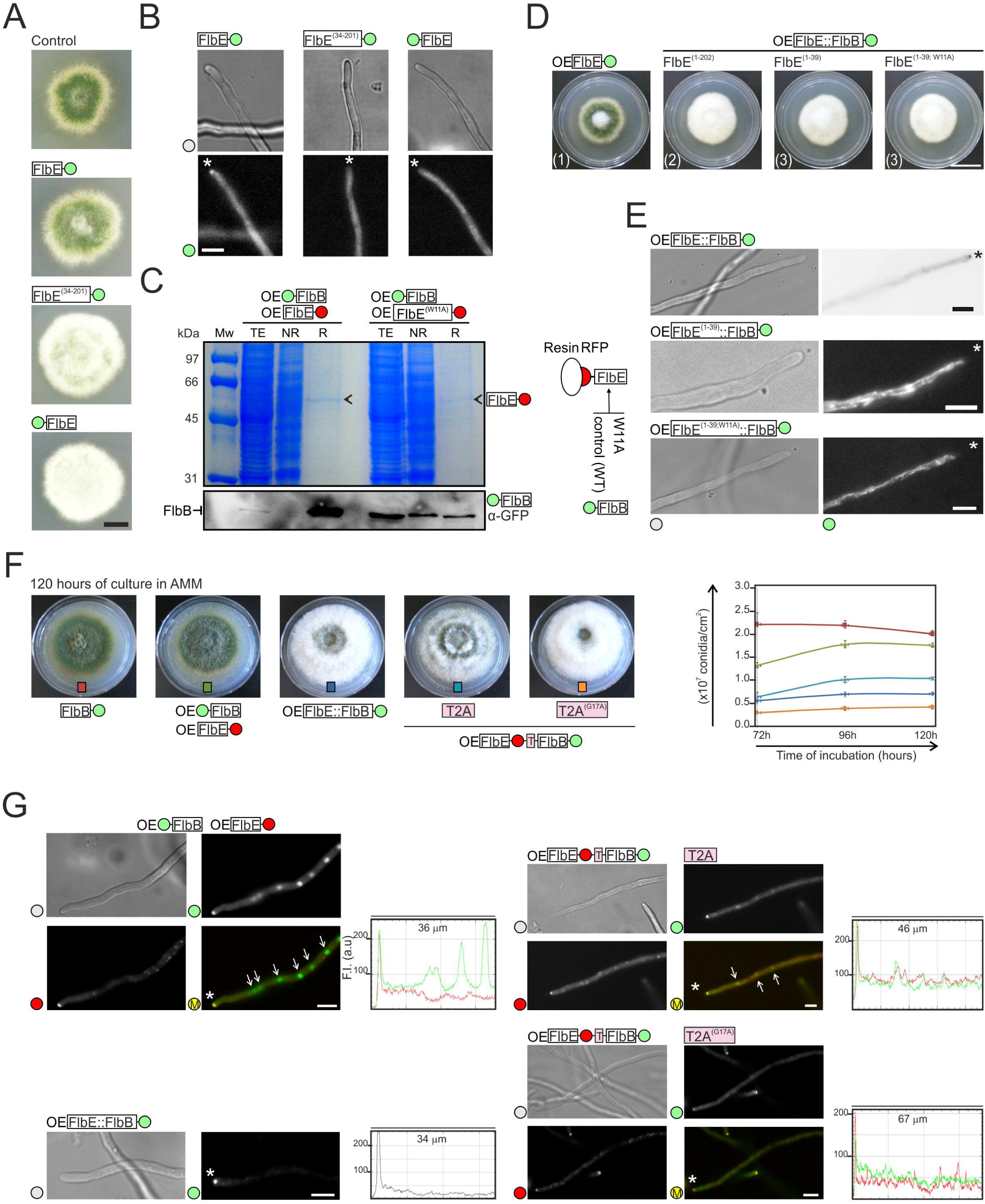
Analysis of FlbE::FlbB chimeras. A) Phenotype of wild-type (control), FlbE::GFP, FlbE^(34-201)^::GFP and GFP::FlbE strains in adequately supplemented AMM plates after 48 hours of culture at 37 °C. Scale bar = 1 cm. B) Subcellular localization of FlbE::GFP, FlbE^(34-201)^::GFP and GFP::FlbE chimeras in vegetative hyphae. Asterisks indicate tips. Scale bar = 5 µm. C) Immunoprecipitation assay using as baits *gpdA*^*mini*^-driven endogenous FlbE::mRFP (left) or FlbE^(W11A)^::mRFP (right) chimeras. The Western-blot below shows the levels of GFP::FlbB in total extracts as well as non-retained (NR) and retained (R) fractions. A Polyacrilamide gel stained with Coomassie blue is shown as control. D) Phenotype of strains expressing *gpdA*^*mini*^-driven (OE) FlbE::GFP (reference), FlbE^(1-202)^::FlbB::GFP, FlbE^(1-39)^::FlbB::GFP or FlbE^(1-39;W11A)^::FlbB::GFP chimeras after 72 hours of culture at 37 °C in MMA. Scale bar = 2 cm. E) Subcellular localization of the above-mentioned chimeras in vegetative hyphae. Asterisks indicate tips. Scale bar = 5 µm. F) Phenotype of strains expressing T2A peptide-containing FlbB and FlbE chimeras (wild-type or the G17A mutant) after 120 hours of culture in plates (diameter = 5.5cm) filled with AMM. Strains expressing 1) FlbB::GFP, 2) FlbE::mRFP and GFP::FlbB, or 3) FlbE::FlbB::GFP chimeras were used as controls. The graph on the right shows conidia production (conidia/cm^2^) for the strains on the left. Values for each strain and time-point are given as the mean of three replicates plus s.e.m. G) Fluorescence microscopy images corresponding to hyphae of strains expressing 1) GFP::FlbB and FlbE::mRFP, 2) FlbE::FlbB::GFP, 3) FlbE::mRFP::T2A::FlbB::GFP and 4) FlbE::mRFP::T2A^(G17A)^::FlbB::GFP chimeras. White asterisks indicate hyphal tips and arrows nuclei. The graphs on the right of each group of images show red and green fluorescence intensities (arbitrary units) in hyphal segments covering the indicated length. Scale bars = 5 µm. See also Figure EV2

Mislocalization of FlbE with a N-terminal GFP tag may be related to failure of any attempt to show an interaction between a bacterially expressed GST::FlbE chimera (used as bait) and FlbB::HA_3x_ (Figure EV2C). Behavior of this type of construct could be due to interaction between the GFP moiety and parts of the FlbE sequence or its interference with localization motifs. However, an immunoprecipitation experiment performed with FlbE::mRFP shows that it can be used successfully as bait, retaining GFP::FlbB when it is in the wild-type form but not when it bears the W11A mutation within domain E1 (Figure 2C).

In order to determine whether domain E1 is sufficient to target FlbB to the tip of hyphae, three DNA constructs were generated. One of them contained the entire FlbE protein tagged in the C-terminus with an FlbB::GFP chimera (Figure EV2D). Second and third constructs bore only the putative signal peptide of FlbE (amino acids from 1 to 39) fused to FlbB::GFP, but the third one included the mutation W11A (FlbE^(1-39; W11A)^::FlbB::GFP) (Figure EV2E). All of them were driven by *gpdA*^*mini*^. Protoplasts of a Δ*flbB* strain were transformed and recombination of DNA constructs was selected at the *flbE locus*, so as to guarantee that the only source of FlbB and FlbE was the one derived from the translation of the constructs. Correct recombination of the constructs was confirmed by Southern-blot and the correctness of the reading frame in the transition from *flbE* to *flbB* sequences was confirmed by genomic sequencing. All transformants showed the characteristic aconidial phenotype of the null *flbE* strain (Figure 2D). Protein chimeras of the expected size were detected in all strains by immunodetection (Figure EV2F).

The fluorescence of the FlbE::FlbB::GFP chimera was detected at the tip of hyphae, suggesting that it can meet all requirement for utilization of the acropetal transport pathway (Figure 2E). Despite the constitutive overexpression provided by the *gpdA*^*mini*^ promoter in this chimera, it was not detected in nuclei (n = 45 hyphae). Considering the *fluffy* phenotype of the strain, it can be hypothesized that FlbB cannot be released from the tip, inhibiting its basipetal transport and thus the transcriptional control of *brlA* in nuclei. In contrast, chimeras bearing only domain E1 of FlbE accumulated in cytoplasmic filamentous structures (Figure 2E) which resembled mitochondria [38]. These results showed that domain E1 of FlbE is not sufficient to target FlbB to the tip and highlighted the importance of additional regions of FlbE for apical localization, such as domains E4, E5, the linker region (see above; [34]), and even domain E3.

To further investigate the hypothetic requirement of an inhibition of the interaction between FlbB and FlbE in order to initiate basipetal transport of the former, additional FlbE chimeras were generated and expressed in a Δ*flbB* background. Constructs were integrated in the *flbE locus* (as above). Transformant strains expressed a *gpdA*^*mini*^-driven chimera consisting of FlbE::mRFP fused to and in frame with FlbB::GFP through a wild-type or a mutant short sequence corresponding to the viral peptide T2A (EGRGSLLTCGDVEENPGP or EGRGSLLTCGDVEENP<u>A</u>P, respectively) (Figure EV2G). During translation of the mRNA, T2A induces the cleavage of the peptide at its last codon (G17-P18) but without blocking translation, which continues until the stop codon of the construct [39]. The maintenance of the correct reading frame was confirmed by sequencing and the synthesis of chimeras of the expected size by immunodetection (Figure EV2H). Peptides with sizes corresponding to FlbE::mRFP and FlbB::GFP were detected in strains expressing the wild-type T2A sequence. However, bands corresponding to the uncleaved, full-length chimera were also detected, as in the case of strains bearing the mutant T2A^(G17A)^ sequence. This suggested that the efficiency of T2A in our system was partial. Accordingly, the strains with the wild-type T2A sequence partially restored conidia production to levels between those of wild-type and aconidial reference strains (Figure 2F). Those results correlated with the subcellular localization of FlbB. When the wild-type T2A peptide was expressed, FlbB recovered nuclear localization but in the form of a weak gradient, despite being expressed driven by *gpdA*^*mini*^ (Figure 2G). In this case, the fluorescence intensity ratio between the tip and the most apical nucleus decreased from 2.78 ± 0.65 in strains expressing the mutant T2A^(G17A)^ form (cytoplasmic fluorescence was considered as the value for nuclei) to 1.60 ± 0.48 when the wild-type T2A form was expressed (p = 0.0000079; n = 15 hyphae for each strain). Overall, results in this section extend our previous model showing that the apical interaction between FlbB and FlbE is in all probability inhibited in order to initiate the basipetal transport of the TF.

### Apical accumulation of FlbB and FlbE, but not their interaction, requires cysteines 272 and 382 of FlbB

The central and C-terminal domains of FlbB are essential for its accumulation at the tip and the induction of conidiation under standard culture conditions [23]. FlbB contains six Cys residues within these domains: Cys236, 272, 280, 303, 382 and 397 (Figure 3A). These Cys residues show higher or lower conservation within FlbB orthologs [35]. Previous results showed that the substitution of Cys382 by an alanine, but not that of Cys397, inhibits conidiation and the apical accumulation of both FlbB and FlbE [23].

**Figure 3:**
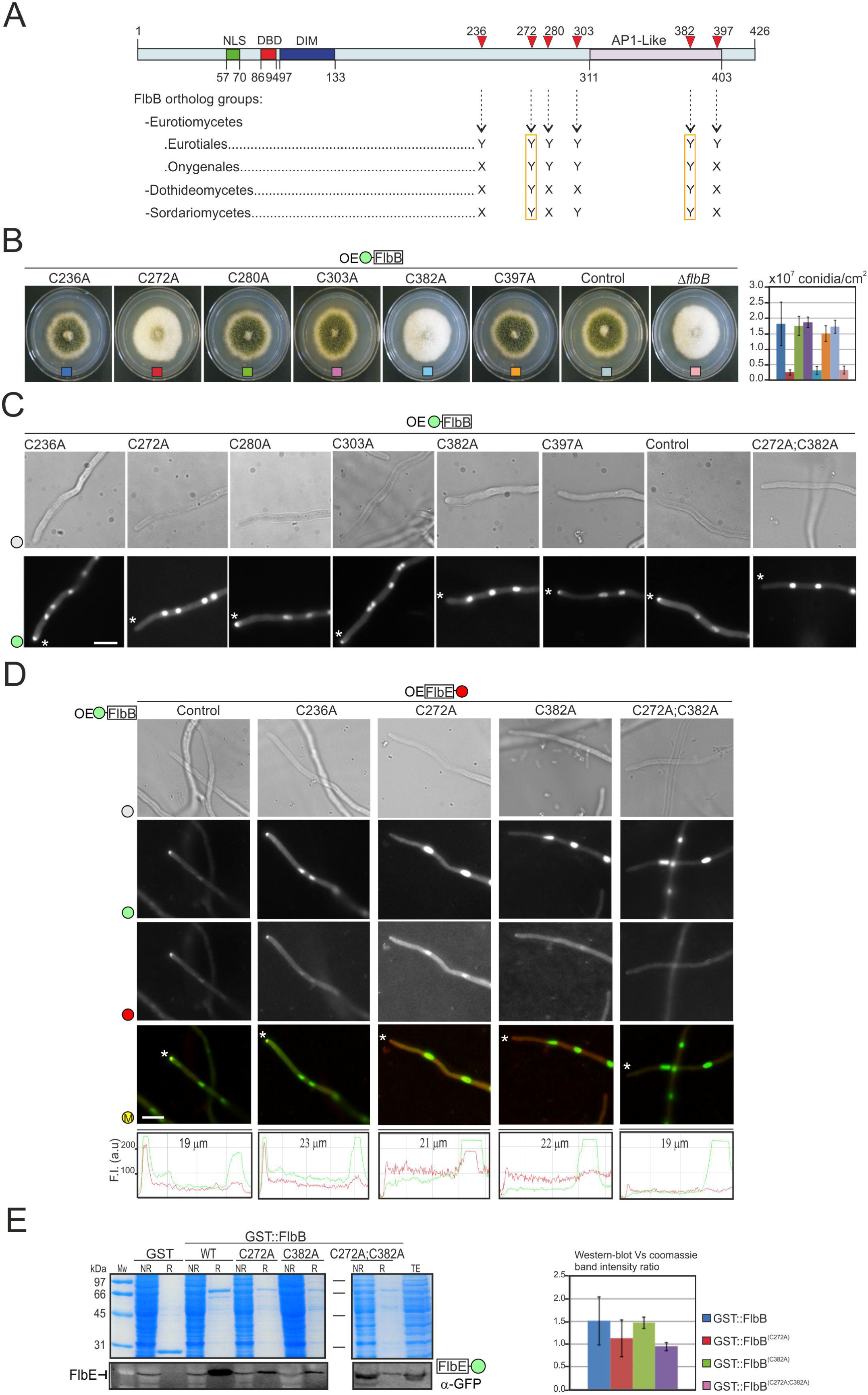
Role of FlbB Cysteines in the apical localization of FlbB and FlbE. A) Diagram showing the position and conservation of the six cysteines within FlbB. “Y” indicates conservation while “X” indicates no conservation. Based on Cortese et al., 2011 [35]. B) Phenotype of the strains expressing *gpdA*^*mini*^ (OE)-driven GFP::FlbB Cys-to-Ala mutant chimeras, on AMM plates (diameter: 5.5 cm) after 72 hours of culture at 37 °C. The graph on the right quantifies conidia produced by each strain (per cm^2^), as the mean of three replicates plus s.e.m. C) Subcellular localization of *gpdA*^*mini*^-driven mutant GFP::FlbB^(Cys-to-Ala)^ chimeras in vegetative hyphae. Tips are indicated by asterisks. Scale bar = 5 µm. D) Subcellular localization of GFP::FlbB and FlbE::mRFP in those FlbB^(Cys-to-Ala)^ mutants showing a *fluffy* phenotype (Cys272Ala. Cys382Ala, and Cys272Ala;Cys382Ala). The reference wild-type strain and the Cys236Ala mutant were used as controls. The graphs at the bottom correspond to hyphae indicated by an asterisk and show green and red fluorescence intensity (arbitrary units) along hyphal segments covering the tip and the most apical nucleus. Scale bar = 5 µm. E) Co-immunoprecipitation assays between bacterially-expressed GST::FlbB forms (wild-type and those Cys-to-Ala mutants causing a *fluffy* phenotype) and crude protein extracts from a strain expressing FlbE::GFP. NR: Not-retained fraction. R: Retained fraction. The graph on the right shows the intensity ratios between western-blot and coomassie bands for each assay. The values are the mean of three replicates plus s.e.m. See also Figure EV3.

In order to assess the importance of these six cysteines in FlbB and FlbE dynamics and functionality, we followed the mutagenesis procedure described in Figure EV3A to construct a set of mutants where each was substituted with an alanine. All mutant constructs were driven by the *gpdA*^*mini*^ promoter. As can be seen in Figure 3B, only the alanine substitutions of Cys272 and Cys382 inhibited conidiation and caused a *fluffy* phenotype as observed previously with the Cys382Ala mutant or the null *flbB* strain. Moreover, conidia production was decreased in only these two strains (2.5-3.2 x 10^6^ ± 0.9-1.3 x 10^6^ conidia/cm^2^ in Cys272Ala, Cys382Ala and null *flbB* strains; n = 3 for each strain; p = 0.00034, 0.00055 and 0.00056, respectively, in the comparison of *fluffy* strains with the reference wild-type strain) compared to the wild-type level of production observed in strains expressing wild-type, Cys236Ala, Cys280Ala, Cys303Ala and Cys397Ala GFP::FlbB chimeras (1.5-1.9 x 10^7^ ± 0.2-0.6 x 10^7^ conidia/cm^2^) (Figure 3B). All chimeras could be detected by Western-blot and showed the same electrophoretic mobility (Figure EV3B; strains integrating one or two copies of the mutant plasmids were analyzed). The aconidial phenotype of those Cys-to-Ala mutants correlated with the absence of FlbB and FlbE from the tip (Figure 3C and 3D). Immunoprecipitation experiments between bacterially expressed wild-type or Cys-to-Ala mutant GST::FlbB forms (Cys272Ala; Cys382Ala or the double mutant Cys272Ala;Cys382Ala) and crude protein extracts of a strain expressing FlbE::GFP (Figure 3E) suggested that these two Cys residues are not essential for the interaction between these two UDA-s. Thus, the de-localization of FlbB/FlbE from the tip observed in those mutants was due to other reasons. These results and those showed in previous sections suggest that there is an inter-dependence between FlbB and FlbE for their transport to and accumulation at the tip, and that the incorporation of the complex to the corresponding transport pathway is enabled by specific residues/domains of both proteins.

### Apical localization of FlbB is altered in a Δ*myoE* background

In cells treated with latrunculin B, which prevents actin monomers from polymerizing, a *gpdA*^*mini*^-driven GFP::FlbB chimera accumulated in the hyphal subapex but was excluded from the apex ([23]; see also Figure 4A). This meant that, under those conditions, FlbB could move in an acropetal direction and reach the subapex but failed to progress to the apex. Those results suggested that the final stage of the acropetal transport of FlbB was dependent on actin filaments while the transport to the subapex was not. *A. nidulans* myosin V, MyoE, has been proposed to fuel the actin filament-dependent step of exocytosis [40,41]. Thus, we generated a Δ*myoE* mutant that expressed an *flbB*^*p*^-driven GFP::FlbB chimera and observed that, compared to the wild-type background, FlbB could not gather at the apex and spread into an apical crescent (Figure 4B). These results suggest that the transport of FlbB from the subapex to the apex occurs through actin filaments and depends on the motor MyoE (see Discussion).

**Figure 4:**
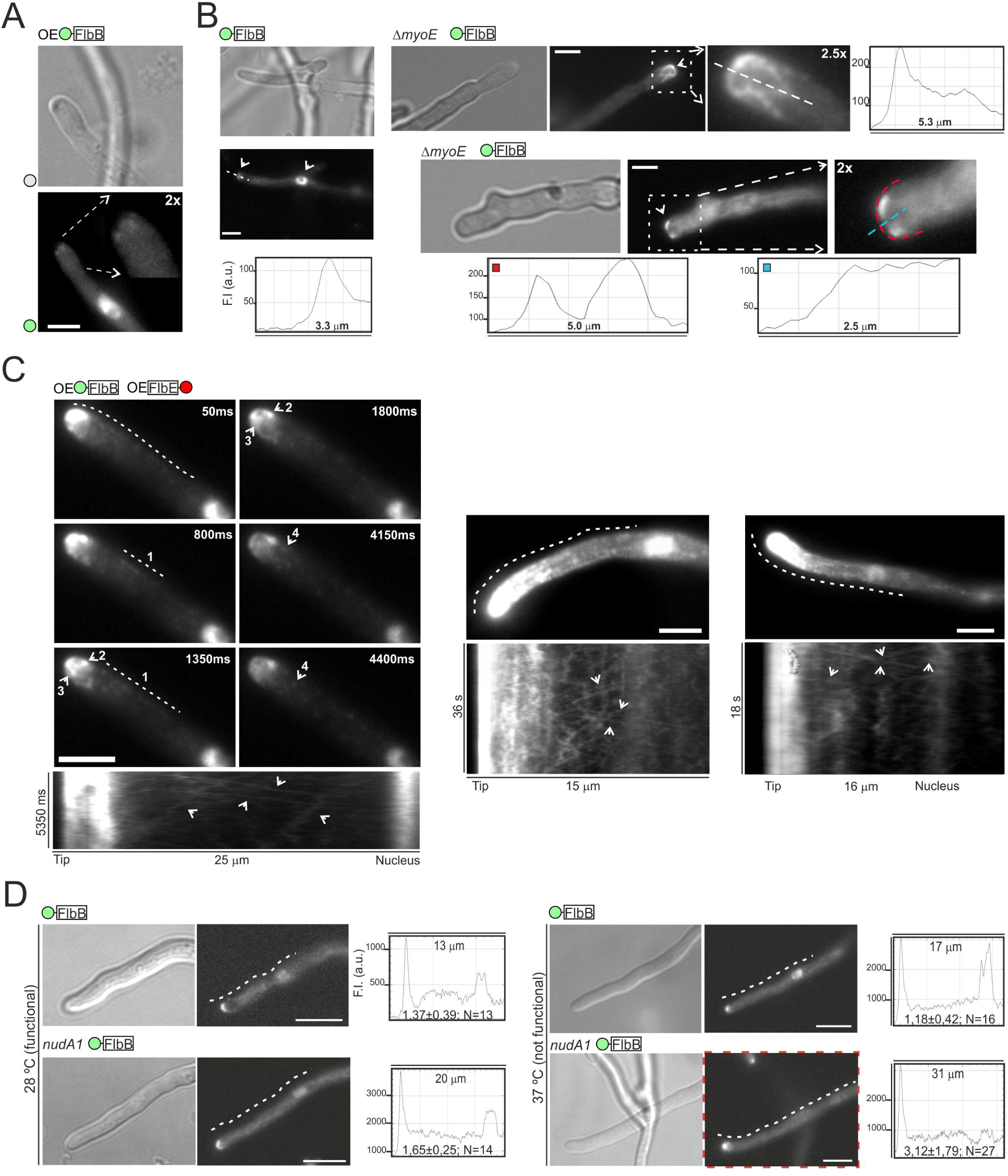
MyoE and NudA in the subcellular localization of FlbB: A) Subcellular localization of a *gpdA*^*mini*^-driven GFP::FlbB chimera in a medium containing Latrunculin B (100 µm) as an inhibitor of actin filament polymerization. See reference [23]. B) Tip localization of GFP::FlbB in wild-type and Δ*myoE*^*myoV*^ genetic backgrounds. The deletion of *myoE* causes the spreading of GFP::FlbB into an apical crescent instead of accumulating at the apex [41]. Microfilament-like structures decorated with GFP::FlbB were also observed. The graphs show the intensity of green fluorescence along the segments indicated by the red and blue dotted lines. In all panels, scale bar = 5 µm. C) GFP::FlbB dynamics in three vegetative hyphae of strains co-expressing FlbE::RFP, both under the control of *gpdA*^*mini*^ (OE). The six frames shown in the left group of images were selected from Video EV4A. The dotted line in the first frame indicates the region analyzed in the kymograph below, which shows patches of GFP::FlbB moving in acropetal and basipetal directions between the tip and the most apical nucleus (number 1). Number 2 indicates a subapical, motionless spot of GFP::FlbB from which basipetal patches depart in frames corresponding to 4150 and 4400 ms (number 4). Finally, number 3 indicates filamentous structures that apparently link the subapex and the apex. The remaining two hyphae (middle and right) correspond to Videos EV4B and EV4C. Scale bars = 5 µm. D) Subcellular localization of GFP::FlbB in wild-type and *nudA1* genetic backgrounds at 28 or 37 °C. The graphs show the intensity of green fluorescence along the hyphal segments covered by the dotted lines. The ratios between the intensity of fluorescence at the tip and the most apical nucleus are also included. The dotted red square indicates the loss of nuclear localization of FlbB when NudA activity is inhibited (*nudA1* background at 37 °C). Scale bars = 5 µm. See also Video EV4.

### Nuclear accumulation of FlbB is inhibited in a *nudA1* background

In order to obtain additional information about the dynamics of FlbB at the hyphal tip, we took advantage of the higher apical accumulation of FlbB in the dual-OE strain expressing GFP::FlbB and FlbE::RFP both under the control of *gpdA*^*mini*^ (see the images and kymographs in Figure 4C; and videos EV4A-C). Being that the apical concentration of the bZIP was significantly higher in that strain compared to a *gpdA*^*mini*^::GFP::FlbB strain (see Figure 1A), we expected that this would enable us to track the movements of FlbB more clearly.

Patches moving in both directions were indeed detected (numbers 1 and 4 in Figure 4C, left; see also arrowheads in the three kymographs shown and videos EV4B and EV4C). A motionless GFP::FlbB spot could be clearly detected at the subapex in Figure 4C, left (number 2), from which the basipetally moving patches departed (Video EV4A). As the fluorescence intensity decreased as a result of the long exposure times, filament-like fluorescent structures could be observed between the subapex and the apex, which could correspond to GFP::FlbB-decorated actin filaments (number 3 in Figure 4C, left; Video EV4A).

Since the FlbB patches moving towards nuclei seemed to depart from the subapical region corresponding to the dynein loading zone [33], we decided to analyze the nuclear localization of a GFP::FlbB chimera (driven by the native promoter) in a strain expressing the NudA1 thermo-sensitive form of NudA, the heavy chain of dynein [42]. When wild-type and *nudA1* backgrounds were compared at 28 °C (functional NudA1), there was an slight increase in the ratio between the fluorescence intensity at the tip and the most apical nucleus, from 1.37 ± 0.39 in the reference GFP::FlbB strain to 1.65 ± 0.25 in the *nudA1* background (n = 12 and 14 hyphae, respectively; p = 0.04; Figure 4D, left). At the restrictive temperature of 37 °C [43], fluorescent nuclei were hardly detected in the *nudA1* background (red square in Figure 4D). The fluorescence intensity ratio between the tip and the most apical nucleus significantly increased from 1.18 ± 0.42 in the wild-type to 3.12 ± 1.79 in the *nudA1* background (n = 16 and 27, respectively; p = 0.00013; since GFP::FlbB was not excluded from nuclei in the mutant we considered the intensity of cytoplasmic fluorescence in hyphae where nuclei could not be discerned). These results support a model in which basipetal transport of FlbB relies principally on the motor complex dynein and its movement towards the minus ends of MTs.

### FlbD is essential for the nuclear accumulation of FlbB

FlbB has a close functional relationship with the cMyb-type UDA TF FlbD [30]. Both participate in the control of conidiation through cooperative binding to the promoter of *brlA* (*brlA*^*p*^). Furthermore, chromatin immunoprecipitation assays showed that FlbB cannot bind *brlA*^*p*^ in the absence of FlbD [30]. These preliminary results suggest that FlbD plays an important role in the transcriptional activity of FlbB, but it is unknown if the cMyb factor is required for the nuclear accumulation of the bZIP. Consequently, we analyzed the localization of FlbB::GFP, driven by the native promoter, in a Δ*flbD* strain that co-expressed the histone H1, HhoA, fused to mCherry [44], as a marker for the nuclei (Figure 5A, left). A statistically significant inhibition of the nuclear localization of FlbB::GFP was observed in the null *flbD* strain, together with increased GFP fluorescence in the apex. The fluorescence intensity ratio between the tip and the most apical nucleus increased from 1.50 ± 0.40 in the wild-type to 13.90 ± 3.00 in the null *flbD* strain (n = 10 hyphae for each strain; p = 4.41 x 10^-12^; see the graphs in Figure 5A).

**Figure 5:**
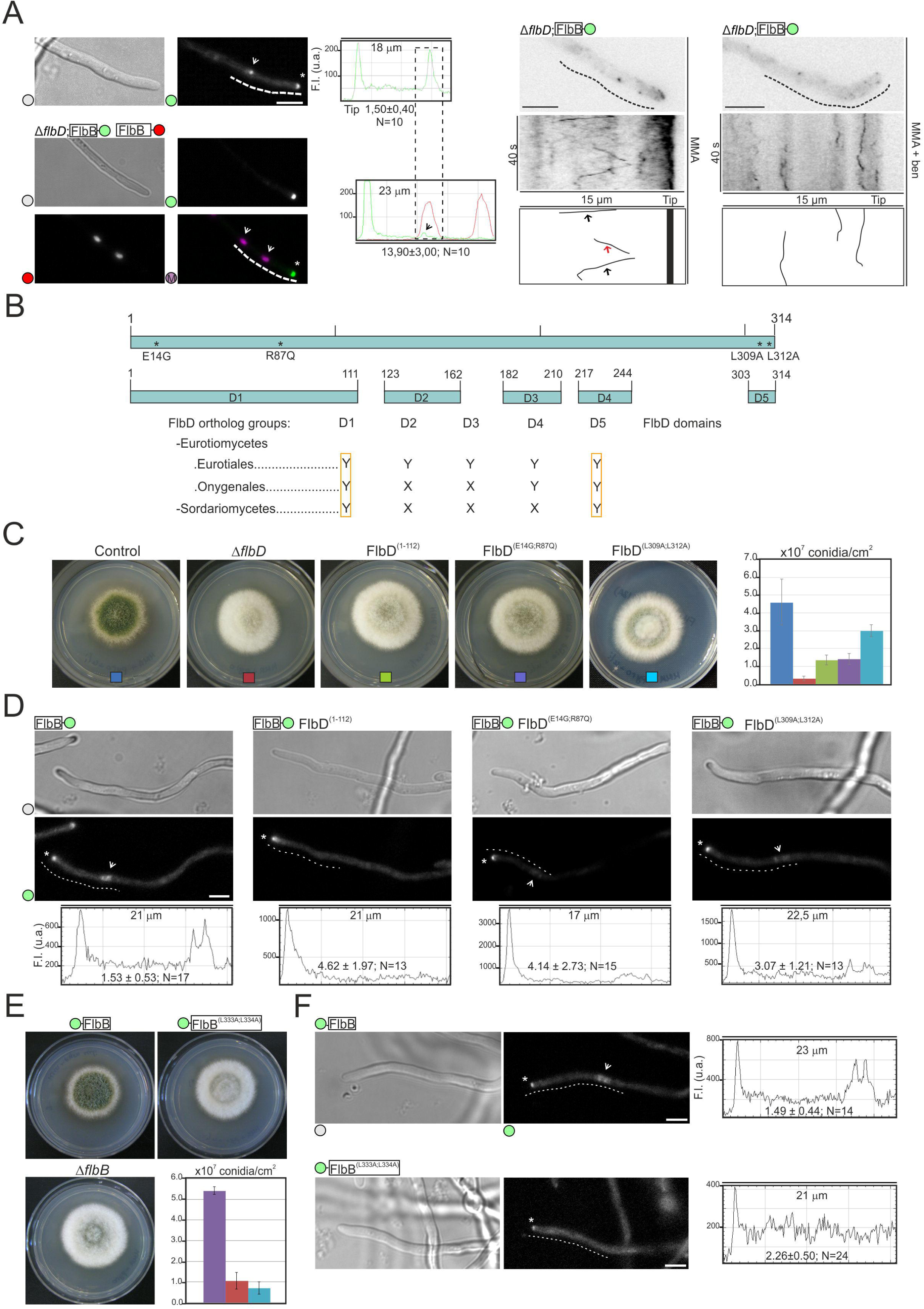
Role of FlbD in the nuclear accumulation of FlbB. A) Left: Subcellular localization of an FlbB::GFP chimera (driven by the native promoter) in vegetative hyphae of wild-type (up) and Δ*flbD* strains (down). In the latter background, nuclei were marked using a HhoA::mCh chimera. Asterisks indicate tips while the arrows indicate nuclei. The graphs on the right show green and red fluorescence intensity (arbitrary units) along the dotted lines. The ratios between the intensity of fluorescence at the tip and the most apical nucleus are also included. Middle: Cytoplasmic movement of FlbB::GFP in a Δ*flbD* genetic background. A hypha with mobile fluorescent FlbB::GFP patches is shown. The kymograph and the diagram below analyze the acropetal (red arrow) and basipetal (black arrows) movement of those patches along the dotted line. Right: Analysis of the dynamics of FlbB::GFP patches in the same strain and in a medium containing benomyl as a MT-destabilizing agent. Vertical lines in the kymograph and the diagram indicate an inhibition of FlbB::GFP movement as result of the addition of the drug. In all panels, scale bar = 5 µm. B) Domain analysis of FlbD sequence. The position and length of each of the five predicted domains, as well as their general conservation in orthologs of specific species within Eurotiomycetes (Eurotiales and Onygenales) and Sordariomycetes, is shown. “Y” indicates conservation while “X” indicates no conservation. The alignment of all the orthologs of FlbD analyzed in this work can be seen in Appendix Figure S1. The position of the mutations within domains D1 (cMyb) and D5 (a predicted LIG_NRBOX) characterized in panels C andD is also indicated. C) Phenotype of wild-type and null *flbD* strains, and strains expressing mutant FlbD^(E14G;R87Q)^, FlbD^(1-112)^ or FlbD^(L309A;L312A)^ forms in AMM plates (diameter = 5.5 cm) after 72 hours of culture. The graph on the right shows conidia production per cm^2^ for each strain at this time-point. Values given are the mean of three replicates plus s.e.m. D) Subcellular localization of GFP::FlbB in the *flbD* mutant backgrounds characterized in panel C. Asterisks indicate hyphal tips and arrowheads, nuclei. Scale bar = 5 µm. E) Phenotype of a strain expressing a GFP::FlbB chimera bearing a double Leu-to-Ala substitution in positions 333 and 334, after 72 hours of culture in AMM and compared to reference GFP::FlbB and Δ*flbB* strains. Diameter of plates is 5.5 cm. The graph shows conidia production per cm^2^ for each strain at the same time-point. Values given are the mean of three replicates plus s.e.m. F) Subcellular localization of wild-type and (L333A; L334A) GFP::FlbB chimeras in vegetative hyphae. Asterisks and the arrowhead indicate hyphal tips and a nucleus, respectively. Scale bar = 5 µm. See also Figure EV5.

Visualizing FlbB movement in vegetative hyphae is difficult. It cannot be detected when FlbB::GFP expression is driven by the native promoter, it can be barely detected near the tip when GFP::FlbB expression is driven by the *gpdA*^*mini*^ promoter [23], and can only be followed when both FlbB and FlbE are expressed constitutively (see above in Figure 4C). Interestingly, deletion of *flbD* allowed the observation of FlbB::GFP patches (*flbB*^*p*^ promoter) moving bidirectionally along the cytoplasm (red and black arrows, respectively; Figure 5A, middle). To check if the bidirectional long-distance cytoplasmic movement of FlbB::GFP in the Δ*flbD* background was MT-dependent, we analyzed FlbB dynamics in a medium containing 3 µg/ml benomyl [45]. The addition of the drug clearly inhibited FlbB::GFP movement, as shown by the vertical lines observed in the kymograph in Figure 5A, right. These results show that in the absence of FlbD, FlbB is not accumulated in nuclei and suggest that it remains moving in both directions along the cytoplasm in a MT-dependent manner.

The above results also suggest that the quantity of FlbB that can be accumulated in nuclei is directly related to FlbD levels. If this hypothesis holds true, overexpression of *flbD* should correlate with a higher nuclear accumulation of the bZIP factor. Thus, *flbD* overexpression was induced through the *alcA* promoter, *alcA*^*p*^ [30]. According to Wieser and coworkers, over-expression of *flbD* triggers the development of conidiophores in shaken cultures, after the transference of mycelia from a liquid medium supplemented with glucose as the carbon source (represses *alcA*^*p*^) to a medium with threonine (*alcA*^*p*^ induction) [46]. This was confirmed for an *alcA*^*p*^::*flbD* strain expressing FlbB::GFP (Figure EV5A).

EtOH (1%) and threonine (100 mM) were assessed as possible carbon sources inducing *alcA*^*p*^-mediated *flbD* overexpression, on solid ACM medium and in comparison with the phenotypes of the reference FlbB::GFP and Δ*flbD* strains (Figure EV5B). In general, all strains produced more aerial hyphae when EtOH was used as the carbon source. The use of threonine, however, induced clear phenotypic differences between *alcA*^*p*^::*flbD* (1.0 x 10^7^ ± 2.2 x 10^6^ conidia/cm^2^) or *alcA*^*p*^::*flbD*; FlbB::GFP (1.2 x 10^7^ ± 1.2 x 10^6^ conidia/cm^2^) and the reference FlbB::GFP strain (5.5 x 10^6^ ± 1.7 x 10^6^ condia/cm^2^; n = 3 for each strain; p = 0.05 and 0.005, respectively).

Considering the results described above, glucose (repressor) or threonine (inducer) were used in the analysis of the subcellular localization of FlbB::GFP in wild-type or *alcA*^*p*^::*flbD* genetic backgrounds (Figure EV5C). Under repressing conditions, FlbB::GFP (wild-type background) localized, as expected, to the tip and the most apical nucleus. The calculated fluorescence intensity ratio between the tip and the most apical nucleus was 1.42 ± 0.39 in this case (n = 10). This ratio increased in the same medium to 4.52 ± 2.05 in the *alcA*^*p*^::*flbD* background (n = 10; p = 0.00012), a significant change that was caused by the decrease in the nuclear localization of FlbB observed when *flbD* expression was repressed (as before, cytoplasmic fluorescence was considered as the value of nuclear fluorescence) (Figure EV5C, upper-right panel). This localization resembled qualitatively what was observed in a Δ*flbD* background (Figure EV5C, bottom-left). Under conditions inducing *alcA*^*p*^, FlbB::GFP recovered the nuclear localization (Figure EV5C, bottom-right), decreasing the fluorescence intensity ratio between the tip and the most apical nucleus to 1.15 ± 0.14 (n = 15; p = 0.000001 compared to the same strain under repressing conditions). Taken together, these results suggest that the *alcA*^*p*^-mediated upregulation of *flbD* in threonine-containing medium increases the nuclear localization of FlbB. Nevertheless, this observation cannot be directly linked to the induction of conidiophore development described in Figure EV5A (shaken cultures) because these fluorescence microscopy analyses were carried out with static instead of shaken cultures. Taken together, the results shown in this section strongly suggest that FlbD is a key element for the nuclear accumulation of FlbB.

### C- and N-termini of FlbD are necessary for conidiation

The role of FlbD in conidiation and the nuclear accumulation of FlbB was then analyzed in more detail. First, we observed that strains expressing N- or C-terminal HA_3x_-tagged versions of FlbD showed a delay in conidiation compared to that expressing an FlbD::GFP chimera (Figure EV5D). Conidia production in HA_3x_-tagged strains was significantly lower than in reference wild-type or FlbD::GFP strains after 48 hours of culture in AMM plates (4.0-4.2 x 10^7^ ± 0.6-1.0 x 10^7^ conidia/cm^2^ in reference strains; 1.0-1.7 x 10^7^ ± 0.2-0.3 x 10^7^ conidia/cm^2^ in HA_3x_-tagged strains; p = 0.65 when the strain expressing a FlbD::GFP chimera and the wild-type strain were compared; p = 0.0000051, 0.00000090 or 0.00000024 when strains expressing FlbD::HA_3x_, HA_3x_::FlbD or HA_3x_::FlbD::GFP chimeras were compared to the reference strain; n = 3 replicates for each strain) (Figure EV5D). These results suggest that HA_3x_ (but not GFP) tagging of FlbD partially hinders its activity.

In an attempt to explain this apparent contradiction (HA_3x_ tag contains nine times less amino acids than GFP), the sequence of FlbD was analyzed. FlbD orthologs were found in Eurotiomycetes (Eurotiales, Onygenales) and Sordariomycetes classes, being the orthologs of this last class the most divergent ones [47]. An alignment of orthologs of Eurotiales differentiated five conserved domains but only two of them, the cMyb transcriptional regulatory domain, which is located at the N-terminus, and a small domain at the C-terminus (residues 303-314), were conserved in all orthologs (Figure 5B and Appendix Figure S1). Thus, we hypothesized that tagging at either the N- or C-termini could partially inhibit FlbD function, delaying conidiation. However, the fact that HA_3x_-tagging but not GFP-tagging delayed conidiation was unexpected.

Since the strain expressing FlbD::GFP conidiated as the wild-type, it could be suggested that the transcript or protein chimera was unstable and therefore truncated, giving an active but untagged version of the protein. A strain expressing an HA_3x_::FlbD::GFP chimera was, thus, generated to confirm this hypothesis through immunodetection experiments (Figure EV5E). Two bands were detected when protein extracts of this strain were hybridized with an α-HA_3x_ antibody, one corresponding to the whole chimera and the second one at a size slightly bigger than that of HA_3x_::FlbD (probably including some amino acids of the N-terminus of GFP). Taken together, these results explain the low fluorescence intensity levels shown by FlbD::GFP [30] and are consistent with both the N-terminal transcriptional regulatory domain and the C-terminal domain playing an important role in FlbD activity.

The study of FlbD forms bearing specific substitutions within the N-terminal region has shown that the cMyb transcriptional regulatory domain is essential to induce or complete both asexual and sexual cycles [48], but there is no information on this region’s hypothetical role in the nuclear accumulation of FlbB. Additionally, the Eukaryotic linear motif (ELM) resource for functional site prediction in proteins (http://elm.eu.org/) suggested that the last C-terminal amino acids of FlbD could correspond to a LIG_NRBOX motif (amino acids 308-314; expect value: 2.63e^-04^), which supposedly confers the ability to bind nuclear receptors and is found primarily in co-activators of those receptors (http://elm.eu.org/elms/LIG_NRBOX.html). Considering the short length of this domain (LxxLL) and the low expect value reported, its presence in FlbD could be meaningless. Thus, we identified all *A. nidulans* proteins predictably containing a LIG_NRBOX domain (2,227 proteins) and observed that transcription factors were significantly enriched in that motif compared to proteins associated to other gene ontology terms (see Appendix Table S1). Therefore, we judged that informatic support of the LxxLL motif of FlbD being functional justified further investigation.

Using a site-directed mutagenesis approach similar to that one followed for *flbE* mutagenesis (Figure EV1C), strains expressing a mutant FlbD^(L309A;L312A)^ form or a truncated FlbD^(1-112)^ form were generated. In addition, using a random mutagenesis approach, an aconidial mutant bearing two point mutations in codons corresponding to the first and second cMyb domains of FlbD (E14G and R87Q) was isolated. The phenotype of these three mutant strains was compared to those of wild-type and null *flbD* strains (Figure 5C). After 72 hours of culture in AMM, conidia production decreased significantly in all mutants compared to the wild-type strain (p = 0.0030, 0.0002 and 0.0152 in strains expressing FlbD^(1-112)^, FlbD^(E14G;R87Q)^ or FlbD^(L309A;L312A)^ forms, respectively; n = 6 for each strain). All these three mutations caused a significant decrease in the ratio between the apical and nuclear fluorescence intensities of FlbB compared to the reference background (p = 0.00056, 0.0000011 and 0.0014, respectively; n > 13 hyphae for each strain) (see Figure 5D), strongly suggesting that besides the DNA-binding domain of FlbD (D1; cMyb), a predicted LIG_NRBOX motif (D5) plays an important role in the nuclear accumulation of the bZIP factor FlbB.

As occurred in FlbD, the presence of a LxxLL sequence was also observed in FlbB (L330 to L334), but not in other TFs known to bind the promoter of *brlA*, such as FlbC, VosA, NsdD or AbaA. The last two Leu residues of this domain were replaced by alanines, FlbB^(L333A;L334A)^, causing a significant decrease in the production of conidia (Figure 5E), from 5.43 x 10^7^ ± 0.2 x 10^7^ in the reference GFP::FlbB strain to 0.71 x 10^7^ ± 0.3 x 10 ^7^ conidia/cm^2^ in the mutant strain (p = 0.0000192; n = 3 for each strain). This phenotype correlated with a significant decrease in the nuclear fluorescence of FlbB. The ratio between the fluorescence intensity at the tip and the most apical nucleus increased from 1.49 ± 0.44 in the reference strain to 2.26 ± 0.50 in the double-leucine mutant of FlbB (p = 0.000029; n = 14 and 24 hyphae, respectively) (Figure 5F). These results support the above-mentioned hypothesis that LxxLL motifs mediate the nuclear localization of UDA TFs FlbB and FlbD.

## Discussion

The activation of the production of asexual multicellular structures in *Aspergillus nidulans* largely (but not exclusively) relies on the signal transduction pathway controlled by FlbB, FlbE and FlbD. A key element for the timely regulation of *brlA* expression is the spatio-temporal control in vegetative hyphae of the dynamics of FlbB, which has to be first transported to the hyphal tip and from there imported into nuclei [23]. Both FlbE and FlbD show a close functional relationship with FlbB and play key roles in this process, but clearly differentiated in space and time. The interaction with FlbE enables acropetal transport and accumulation of FlbB at the tip while FlbD is essential for the nuclear localization of the pool of transcriptionally active FlbB generated at the growth region. The available information and our hypotheses on the molecular basis of this sequence of events has been summarized in Figure 6 and will be used to structure this discussion.

**Figure 6:**
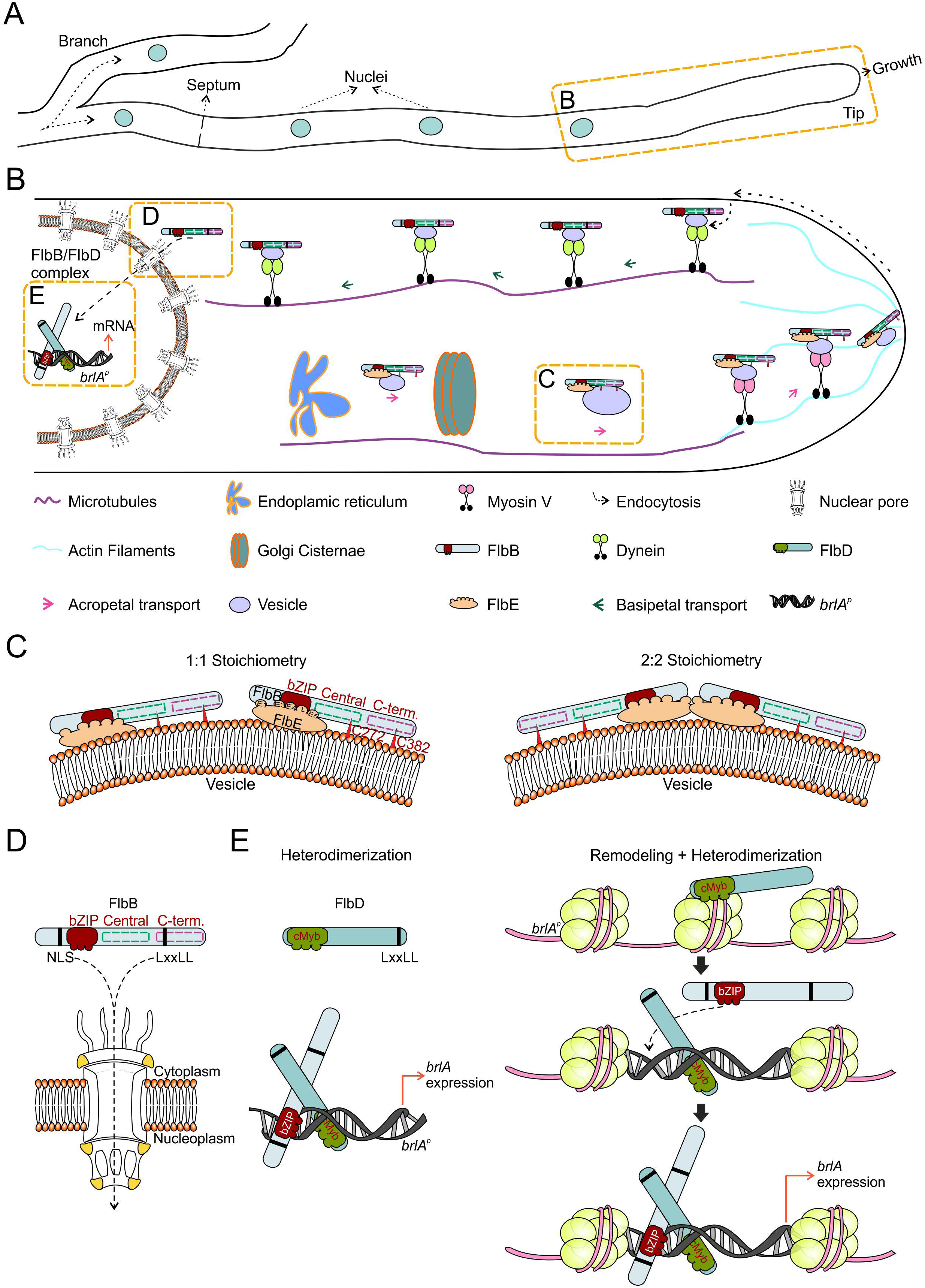
Working model for FlbB dynamics in hyphae of *A. nidulans.* A) Diagram showing the morphology of a hypha, including a branch, a septum (rings that separate cells within hyphae), nuclei and an actively growing tip. B) Magnification of the region between a tip and its closest nucleus, showing a general model for the acropetal and basipetal transport of FlbB. Each player is indicated below. Dotted orange squares mark the features that are analyzed in more detail in the following panels. C) Two hypothetic configurations of the signaling complex formed by FlbB and FlbE, 1:1 (left) or 2:2 (right). In both of them, FlbB interacts with FlbE through the bZIP domain. At least five domains of FlbE (E1, E4, the linker L region, E5 and, to lower extent, E3) would participate in the interaction with the bZIP of FlbB (see also [34]). At least domain E1 would link the complex to the corresponding transport pathway, maybe a vesicle attached to a still unknown molecular motor. Cys382 [23] and Cys272 of FlbB are required for the acropetal transport of the FlbB/FlbE complex but not for their interaction. They could mediate an interaction with the transport vesicle or be required in order to acquire a specific three-dimensional conformation essential to join the transport pathway (this last option is not considered in the model). Finally, myosin V, MyoE, would transport the complex from the subapex to the growing apex of the tip. D) At the hyphal subapex, FlbB would join a dynein-mediated basipetal transport pathway (probably attached to a vesicle or an early-endosome) that would approximate the TF to the nucleus. A NLS and a LxxLL motif are required for the import of FlbB across nuclear pores and its accumulation in nuclei (see also [23]), as well as the cMyb domain and an additional LxxLL motif of FlbD (see panel E). E) Both FlbB and FlbD bind a common region of 300 bp within the promoter of *brlA, brlA*^*p*^ [30]. Based on previous publications describing the interaction of cMyb and bZIP proteins for the control of gene expression [53], a heterodimerization model is proposed on the left. Considering the essentiality of FlbD for the nuclear accumulation of FlbB, as well as the similarities between cMyb (transcriptional regulation) and SANT (chromatin remodeling) domains (http://www.aspergillusgenome.org/cgi-bin/protein/proteinPage.pl?dbid=ASPL0000052812&seq_source=A.%20nidulans%20FGSC%20A4), a more speculative “Remodeling + Heterodimerization” model is proposed. In this model, FlbD binds *brlA*^*p*^ first, inducing, maybe in combination with other proteins such as GcnE or LaeA [54,55], a chromatin remodeling event that enables heterodimerization with FlbB and binding of both TFs to their targets at *brlA*^p^. The NLS of FlbB improves DNA binding [59] and the LxxLL motifs of both FlbD and FlbB may play a modulatory role.

### Acropetal transport mechanism

The results shown in this and previous works demonstrate that the levels of FlbE are directly related to the quantity of FlbB accumulated at the tip. The nuclear, non-transcriptionally active pool of FlbB described when only FlbB is expressed constitutively decreases as the quantity of FlbE increases and disappears in those strains expressing FlbE and FlbB fused in the same chimera (*gpdA*^*mini*^-driven FlbE::FlbB::GFP or FlbE::mRFP::T2A^(G17A)^::FlbB::GFP). These results suggest that the FlbB/FlbE complex is composed of equimolar amounts of each developmental regulator, although the stoichiometry of the complex (i.e., 1:1 or 2:2) is still unknown (Figure 6A, 6B and 6C). While the bZIP domain of FlbB is essential and sufficient for this heterodimerization and to date FlbB homodimers have not been detected [23], cysteine residues in positions 272 and 382 apparently play a modulatory role (Figure 6C). However, that these cysteines are essential for the apical accumulation of the complex, strongly suggests that FlbB does not join its acropetal transport pathway exclusively through FlbE and that its cysteines, probably in combination with additional elements, play a key role (see the legend of Figure 6C).

The role of FlbE in the subcellular dynamics of FlbB seems to be limited exclusively to acropetal transport and at least five of its seven domains (E1, E4, E5, the linker domain and to lesser extent E3) [34] are necessary for the apical accumulation of the bZIP TF (Figure 6B and 6C). Considering that FlbE interacts with the bZIP domain of FlbB (but apparently not with central and C-terminal domains) and the presence of a nuclear localization signal (NLS) prior to the bZIP [23], it is tempting to suggest that besides assisting the acropetal transport of the complex, FlbE binding could occlude the NLS of FlbB, precluding its basipetal transport and nuclear import.

The characteristics of FlbB transport towards the polarity site as well as the possibility of domain E1 of FlbE being a signal peptide open the possibility of the incorporation of the FlbB/FlbE complex into a vesicular fraction that would transit through the ER-Golgi network (Figure 6B). In preliminary LC-MS/MS-coupled co-immunoprecipitation assays of protein extracts of a strain expressing the *gpdA*^*mini*^-driven GFP::FlbB chimera, we identified several proteins participating in the transport between the ER and the Golgi apparatus. These preliminary results correlate with the hypothesis proposed above and at the same time open an avenue for a future, comprehensive analysis of how these two developmental regulators join the secretory pathway, which additional proteins they establish interactions with, which of their domains are required or what could be the conformation and stoichiometry of the complex. The transit of vesicles between the ER and the Golgi apparatus is MT dependent, while FlbB reaches the apex of hyphae in a culture medium containing benomyl, which destabilizes MTs [23]. Thus, additional experiments are required to elucidate the hypothetic mechanism of FlbB/FlbE transition through the ER-Golgi network.

In the absence of actin polymerization, FlbB reaches the subapex but fails to accumulate in the apex [23]. In a null *myoE* background, FlbB spreads into an apical crescent that resembles the localization shown in that genetic background by the post Golgi-carrier marker RabE/Rab11 [41,43]. As Pantazopoulou and collaborators found in their characterization of RabE, it could be suggested that, without MyoE activity, FlbB might be captured by the “actin mop” but lacked a molecular motor which could deliver it to the apex [41]. Thus, myosin V (MyoE) arises as the best candidate motor protein to deliver FlbB, on actin filaments, from the subapex to the apex (Figure 6B).

### Basipetal transport and nuclear accumulation of FlbB

The results shown here suggest that FlbB departs from the dynein loading region in its journey towards nuclei (Figure 6B). In accordance with the long-distance basipetal transport of vesicles and macromolecular cargo in neurons, it is generally accepted in *A. nidulans* hyphae that vesicles formed by endocytosis at the subapex are bound by the dynein complex and transported on MTs towards distal regions [49]. The inhibition of the nuclear accumulation of FlbB observed in a thermo-sensitive heavy-chain dynein *nudA1* background at the restrictive temperature (37 °C) correlates with a model in which FlbB follows this pathway. This, at the same time, is in agreement with the MT-dependence of the cytoplasmic, bidirectional transport of FlbB observed in a Δ*flbD* background, which suggests that in the absence of FlbD, a fraction of FlbB remains moving bidirectionally along the length of the cytoplasm. An interesting question for the future will be the elucidation of how FlbB joins the dynein-mediated basipetal transport pathway and the identification of adaptor proteins and the karyopherins required for its nuclear accumulation. This will open the possibility of a deeper analysis of the role of the NLS and the LxxLL motif of FlbB, both of them required for the nuclear accumulation of the TF (Figure 6B and 6D).

In the absence of FlbD, FlbB does not accumulate in nuclei and cannot bind *brlA*^*p*^ [30], thus inhibiting conidiation. Besides offering the possibility of using the null *flbD* strain to identify proteins required for the basipetal transport and nuclear accumulation of FlbB, these results raise the following question: which is the primary cause of the inability to trigger asexual development in this background. It is clear that the absence of FlbB from nuclei impedes binding to *brlA*^*p*^ but, at the same time, FlbB accumulation in nuclei may be reduced due to an inability to bind DNA in the absence of FlbD activity (Figure 6B and 6E). The only subcellular localization described for FlbD and the orthologs that have been functionally characterized is nuclear [30,50,51], suggesting that it is not directly involved in the basipetal transport of FlbB but in its nuclear retention. Our results show that both the N- and C-termini of FlbD are necessary for the induction of conidiation and the nuclear accumulation of FlbB. Since its N-terminal cMyb transcriptional regulatory domain is sufficient for FlbD to bind *brlA*^*p*^ [30], a link between DNA-binding by FlbD and the nuclear accumulation of FlbB (and perhaps DNA-binding by the bZIP TF) can be suggested. In this context, the possibility of FlbD acting as a pioneer TF is open [52], binding *brlA*^*p*^ first [30], causing a modification of the conformation of chromatin and enabling then binding of FlbB (Figure 6E) [52,53]. Alternatively (or in addition), the cMyb domain of FlbD could act as a heterodimerization domain for the bZIP of FlbB [53], forming a heterocomplex which, in turn, is capable of binding to the targets of each TF within *brlA*^*p*^, which are predicted to be adjacent (Figure 6E) [30]. In both scenarios, LxxLL motifs of both FlbD and FlbB may play a modulatory role and/or mediate in the interaction with additional elements. Although both TFs bind to a common region of 300 nt-s within *brlA*^*p*^, future experiments must determine the nature of the exact target-DNA sequences of both FlbB and FlbD in *brlA*^*p*^, the hierarchy/democracy between both TFs and the study of hypothetical modifications in the structure of chromatin at this region [54,55]. The extension of these analyses to other activators and repressors that bind *brlA*^*p*^ (Lee et al., 2016) will further a deeper understanding of how TFs belonging to different pathways are coordinated for a timely control of multicellular development.

## Materials and Methods

### Oligonucleotides, strains and culture conditions

Appendix Tables S3 and S4 show, respectively, the oligonucleotides and the strains of *A. nidulans* used in this work. Strains were cultivated in supplemented liquid or solid *Aspergillus* minimal (AMM) or complete (ACM) media [56,57], using glucose (2%) and ammonium tartrate (5 mM) as carbon and nitrogen sources, respectively. Fermentation medium (AFM), which contained 25g/L corn steep liquor (Sigma-Aldrich) and sucrose (0.09M) as the carbon source, was used to culture samples for protein extraction [58]. Mycelia for DNA extraction and Southern-blot analysis were cultured in liquid AMM and the procedures described previously by us were followed [34].

Conidia production on solid medium was calculated as described previously by us [30], using three replicates for each strain. The two-tailed Student’s t-test for unpaired samples was used to determine the statistical significance of the changes in conidia production.

Gene overexpression through the use of *alcA*^p^ was induced in solid medium that contained threonine (100 mM) and repressed when glucose (2%) was used. For the *alcA*^*p*^-mediated overexpression in liquid culture, first 10^6^ conidia per milliliter were cultured at 37 °C for 18 hours in standard AMM. Then, mycelia samples were filtered and transferred to AMM that contained threonine (100mM) as the carbon source, with additional 20 hours of culture [59]. Hyphal and conidiophore morphology were then analyzed using a Nikon Optiphot microscope, coupled to a Nikon FX-35DX camera.

Fluorescence microscopy analyses were conducted by inoculating conidiospores of *A. nidulans* strains in supplemented watch minimal medium (WMM) [60] and incubating them for 18 hours at room temperature.

As a sample and a tutorial of the multiple advantages for genetic manipulation offered by *A. nidulans*, the procedures followed for the generation of strains expressing wild-type or mutant forms of the proteins of interest, expressed through native or constitutive promoters, and tagged at N- or C-termini are briefly described along the results section as expanded view figures. Most of those procedures, as well as those followed for the generation of deletants, are based on the fusion-PCR technique developed by Yang and colleagues and the subsequent protoplast transformation protocol developed by Tilburn and colleagues or Szewczyk and colleagues [61–63]. Cys-to-Ala mutants of FlbB were generated by transforming protoplasts with mutant p*gpdA*^*mini*^::GFP::FlbB^(Cys-to-Ala)^ plasmids. Recombination of these mutant plasmids was directed to the *pyroA locus*. The strain expressing GFP::FlbB in a *nudA1* genetic background was generated by meiotic crosses [64].

### Fluorescence microscopy

Subcelllar localization of FlbB and FlbE in hyphae was analyzed as previously described using a Leica DMI-6000b or a Zeiss Axio Observer Z1 inverted microscopes [23,65]. The former is equipped with a 63x Plan Apo 1.4 N.A. oil immersion lens from Leica, and filters GFP (excitation at 470 nm and emission at 525 nm) and Txred (excitation at 562 nm and emission at 624 nm). The latter includes a 63x Plan Apochromat 1.4 oil immersion lens, and filters 38 (excitation at 470 nm and emission at 525 nm) and 43 (excitation at 545 nm and emission at 605 nm). Fluorescence levels were measured using ImageJ software (http://imagej.nih.gov/ij) (U. S. National Institutes of Health, Bethesda, Maryland, USA).

### Protein extraction and immunodetection

Two different protocols were used for protein extraction. Direct immunodetection of proteins was done using protein extracts that were obtained through the alkaline lysis protocol [66], which prevents protein degradation. Briefly, approximately 6 mg of lyophilized mycelium were resuspended in 1 ml lysis buffer (0.2M NaOH, 0.2 % β-mercaptoethanol). After trichloroacetid acid (TCA) precipitation, 100 µl Tris-Base (1 M) and 200 µl of loading buffer (62.5 mM Tris-HCl pH=6.8, 2 % SDS (p/v), 5 % β-mercaptoethanol (v/v), 6 M urea and 0.05 % bromophenol blue (p/v)) were added. Samples were then loaded on polyacrilamide gels (%10) for protein-content assessment and Western-blot analysis.

For co-immunoprecipitation assays, protein extracts were obtained through the procedure described by [67]. Samples were lyophilized and pulverized before the addition of 1 ml of NP-40 extraction buffer (5 mM Hepes pH=7.5, 1 mM EDTA, 20 mM KCl, 0.1% NP-40, 0.5 mM DTT and a protease inhibitor cocktail from Roche; plus 150 mM NaCl when the extract was going to be incubated with Chromotek’s GFP-Trap beads). The Bradford assay was used for the determination of protein concentration. Before the use of crude extracts in co-immunoprecipitation assays, 200 µg of protein were precipitated with TCA and purified with ethanol/eter mixes (1:1 and 1:3, respectively). After resuspension in 80 µl of loading-buffer, the integrity of samples was assessed by polyacrilamide gel electrophoresis while expression of the chimera of interest was confirmed by Western-blot.

For immunodetection analyses, proteins were separated in standard 10% SDS-polyacrylamide gels before electro-transferring them to nitrocellulose filters. GFP-, mRFP- or HA_3x_-tagged proteins were detected with α-GFP (mouse), α-RFP (rabbit) or α-HA_3x_ (mouse) monoclonal antibody cocktails (1:5000 Roche, 1:4000 USBiological and 1:1000 Santa Cruz, respectively). Peroxidase conjugated α-mouse or α-rabbit (1/4000, Jackson ImmunoResearch Laboratories, or 1:10000, Sigma Aldrich, respectively) were used as secondary antibody. Peroxidase activity was induced with Amersham Biosciences ECL kit, and chemiluminescence was detected using a Chemidoc +□XRS system (Bio-Rad).

### Expression of recombinant proteins in Bacteria

Plasmid pGEX-2T (Pharmacia) was used to express fusions of GST to full (1-426) or point-mutant (C272A, C382A and C272A;C382A) versions of FlbB. After culturing transformant *E. coli* DH1 cells until OD600□=□0.6–0.8, expression of GST-tagged proteins was induced with the addition of 0.1□mM IPTG and further incubation at 15°C for 24□h. Extracts containing recombinant GST chimeras were then obtained essentially as described by us previously [23].

### Co-immunoprecipitation assays

Two procedures were followed. The first one analyzed the ability of the above-mentioned GST-tagged FlbB chimeras, used as baits, to retain wild-type and mutant forms of FlbE::GFP [23]. Briefly, GST-tagged proteins attached to glutathione sepharose media (GE Healthcare) were incubated at 4°C for 1 h with 2-3 mg of crude protein extracts of *A. nidulans*. After 3-5 washing steps (a sample of this non-retained, NR, fraction was stored for analysis), loading buffer was used to resuspended the resin (retained fraction, R). Proteins were visualized with the Bio-Safe Coomassie stain (Bio-Rad), and tagged proteins were specifically detected by immunodetection.

In the second procedure, 25 µL of GFP- or RFP-Trap beads (Chromotek) were washed twice by centrifugation at 2,500g and 4 °C with 500 µL dilution buffer (5 mM Hepes pH = 7.5, 150 mM NaCl and 0.5 mM EDTA) and incubated with 6 mg of the protein extract for 90 minutes at 4 °C. After centrifugation, the supernatant was precipitated in TCA, resuspended in loading buffer and stored as the non-retained (NR) fraction. The resin was then washed three times using protein extraction buffer plus 150 mM NaCl (see before) and finally resuspended in loading buffer (R: retained fraction). Both NR and R fractions were resolved by SDS-PAGE electrophoresis.

## Supporting information

Main Figure 5

Oligonucleotides used in this work

Strains of A. nidulans used in this work

Main Figure 5

## Author contribution

O.E., E.A.E., A.O., E.P-A. and E.O-A designed and generated the strains, and carried out the experiments. O.E. and E.A.E. supervised the experimental part. O.E. wrote the manuscript. M.S.C. performed bioinformatic analyses. All authors contributed to the improvement of the text and figures.

## Funding information

Work at the UPV/EHU lab was funded by UPV/EHU (grant EHUA15/08 to O.E) and the Basque Government (grant IT599-13 to Dr. Unai Ugalde). Work at CIB-CSIC was funded by MINECO (BFU2015-66806-R to E.A.E). E.P-A and E.O-A held predoctoral fellowships from UPV/EHU. A.O held a predoctoral fellowship from the Basque Government.

### Acknowledgements

We want to acknowledge the work done by Ion Luis Abad, Luis Pablo Gonzalo and Alba Ledesma, all of them former students at UPV/EHU, in the generation of point *flbE* mutants (W11A) and the strains expressing the T2A-tagged forms, as well as the bioinformatic analysis of FlbD.

## Supplementary material

**<u>Appendix Table S1</u>: Enrichment of the LxxLL motif in transcription factors of *A. nidulans***. To analyze if the LxxLL motif is enriched in TFs of *A. nidulans* compared to other GO terms, the work by Wortman and colleagues was taken as a reference [68]. The authors predicted the presence of 490 TF-coding genes in *A. nidulans*, a 4,6 % of a total of 10,701 protein-coding genes. A total of 2,227 *A. nidulans* proteins containing at least one LxxLL motif were identified. FlbB and FlbD were the only known TFs controlling conidiation with such a motif in their amino acid sequences. The table compares the number of LxxLL motif-containing proteins with GO terms associated with TFs, such as Transcription, Nucleus, Nuclear or Transcription factor, with the total number of proteins associated to those GO terms in the *A. nidulans* proteome. The enrichment is calculated as the ratio between these parameters, and compared to other GO terms not directly associated with TFs, such as translation, mitochondria, activity and metabolic.

**<u>Appendix Table S2</u>: Oligonucleotides used in this study.**

**<u>Appendix Table S3</u>: Strains used in this study.**

**<u>Appendix Figure S1</u>**: Sequence alignment of FlbD orthologs, obtained using the Genedoc software (version 2.7.000). Green boxes indicate the extension of the FlbD regions conserved in the orthologs of species from the Eurotiales order. Those domains that are conserved in all orthologs (N- and C-terminal domains) are in red. Nomenclature: Aory, *Aspergillus oryzae*; Afla, *A. flavus*; Anig, *A. niger*; Ater, *A. terreus*; Acla, *A. clavatus*; Afum, *A. fumigatus*, Nfis, *Neosarotya fischeri*; Tsti, *Talaromyces stipitatus*; Ptri, *Pyrenophora tritici*; Sscl, *Sclerotinia sclerotorium*; Tver, *Trichophyton verrucosum*; Aben, *Arthroderma benhamiae*; Trub, *T. rubrum*; Cpos, *Coccidioides posadasii*; Cimm, *C. immitis*; Ader, *Ajellomyces dermatitidis*; Pbra, *Paracoccidioides brasiliensis*; Fpse, *Fusarium pseudograminearum*; Foxy, *F. oxysporum*; Fver, *F. verticillioides*; Vdah, *Verticillium dahliae*; Ndis, *Neurospora discreta*; Smac, *Sordaria macrospora*; Ncra, *N. crassa*; Ntre, *N. tetrasperma*.

**Figure EV1:**
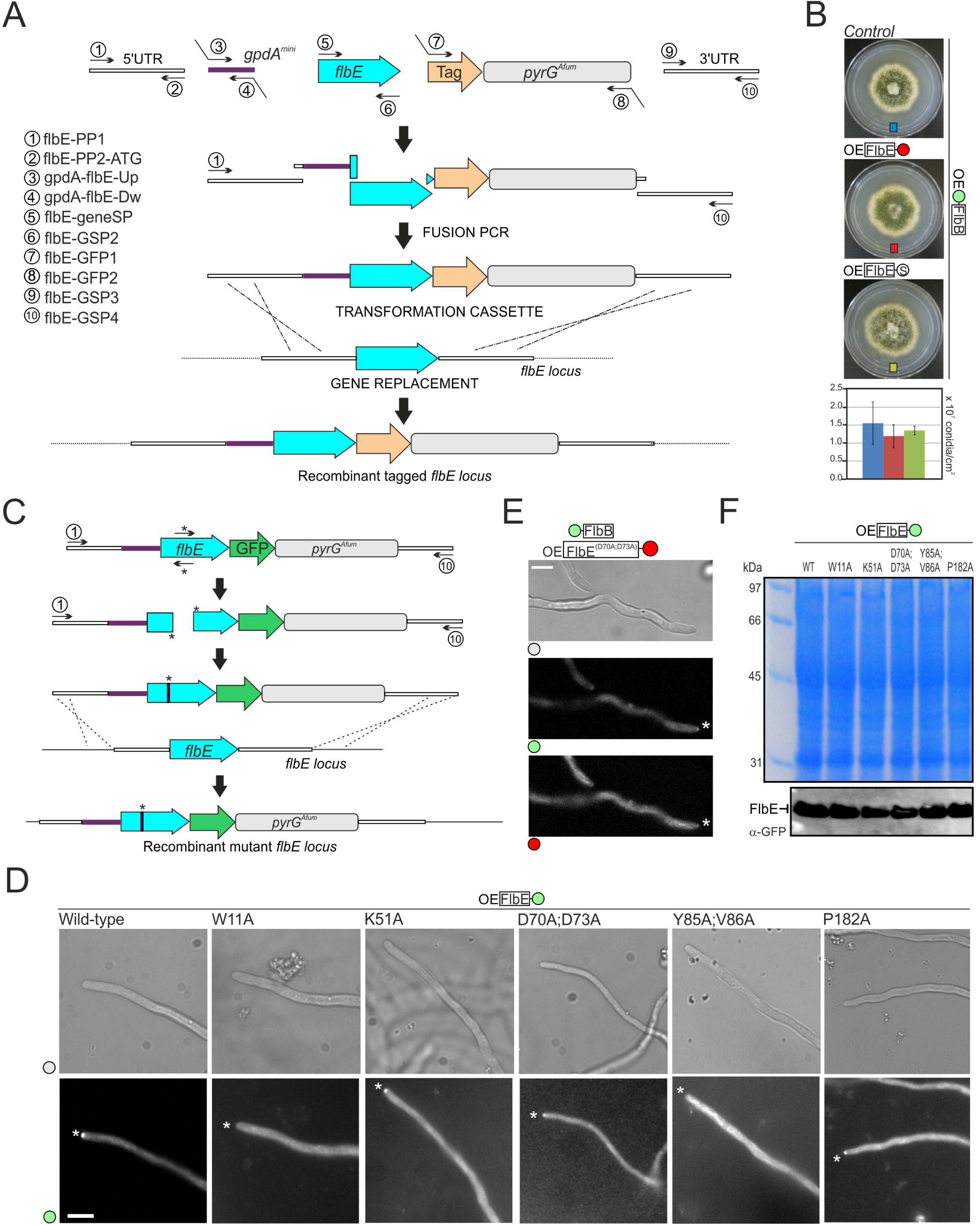
FlbE analysis. A) Procedure followed for the generation of a dual-OE strain co-expressing GFP::FlbB and FlbE::GFP, both driven by *gpdA*^*mini*^. The promoter of *flbE, gpdA*^*mini*^, *flbE* coding region, RFP or Stag plus the *pyrG* gene from *A. fumigatus*, and the 3’-UTR region were amplified independently and fused. The amplicon was used to transform protoplasts of a strain expressing GFP::FlbB driven by *gpdA*^*mini*^. Recombination was induced at the *flbE locus*. B) Phenotype of the strains analyzed in Figure 1A after 72 hours of culture in solid AMM at 37 °C (diameter of plates is 5.5 cm; OE: *gpdA*^*mini*^-driven). The graph below quantifies conidia production for each strain (control: 1.56 x 10^7^ ± 0.6 x 10^7^ conidia / cm^2^; mRFP-tagged strain: 1.2 x 10^7^ ± 0.3 x 10^7^ conidia / cm^2^; S-tagged strain: 1.35 x 10^7^ ± 0.1 x 10^7^ conidia / cm^2^), calculated as the average value of three replicates plus s.e.m. C) Strategy followed for the generation of strains expressing *gpdA*^*mini*^-driven mutant FlbE*::GFP strains. Using oligonucleotides bearing the designed *flbE* mutations, two PCR fragments were synthesized and fused. The fusion construct was used to transform protoplasts of a wild-type, TN02A3 [69], strain. D) Subcellular localization of *gpdA*^*mini*^-driven wild-type and mutant FlbE::GFP chimeras in vegetative hyphae. Asterisks indicate hyphal tips. Scale bar = 5 µm. E) Subcellular localization of GFP::FlbB (driven by the native promoter) in a strain that expresses constitutively a mutant FlbE^(D70A;D73A)^::mRFP chimera. The asterisk indicates the tip. Scale bar = 5 µm. F) Immunodetection of *gpdA*^*mini*^ (OE)-driven wild-type and mutant FlbE::GFP chimeras in crude extracts of the corresponding strains. The Coomassie-stained gel is shown as a loading control. See also Figure 1.

**Figure EV2:**
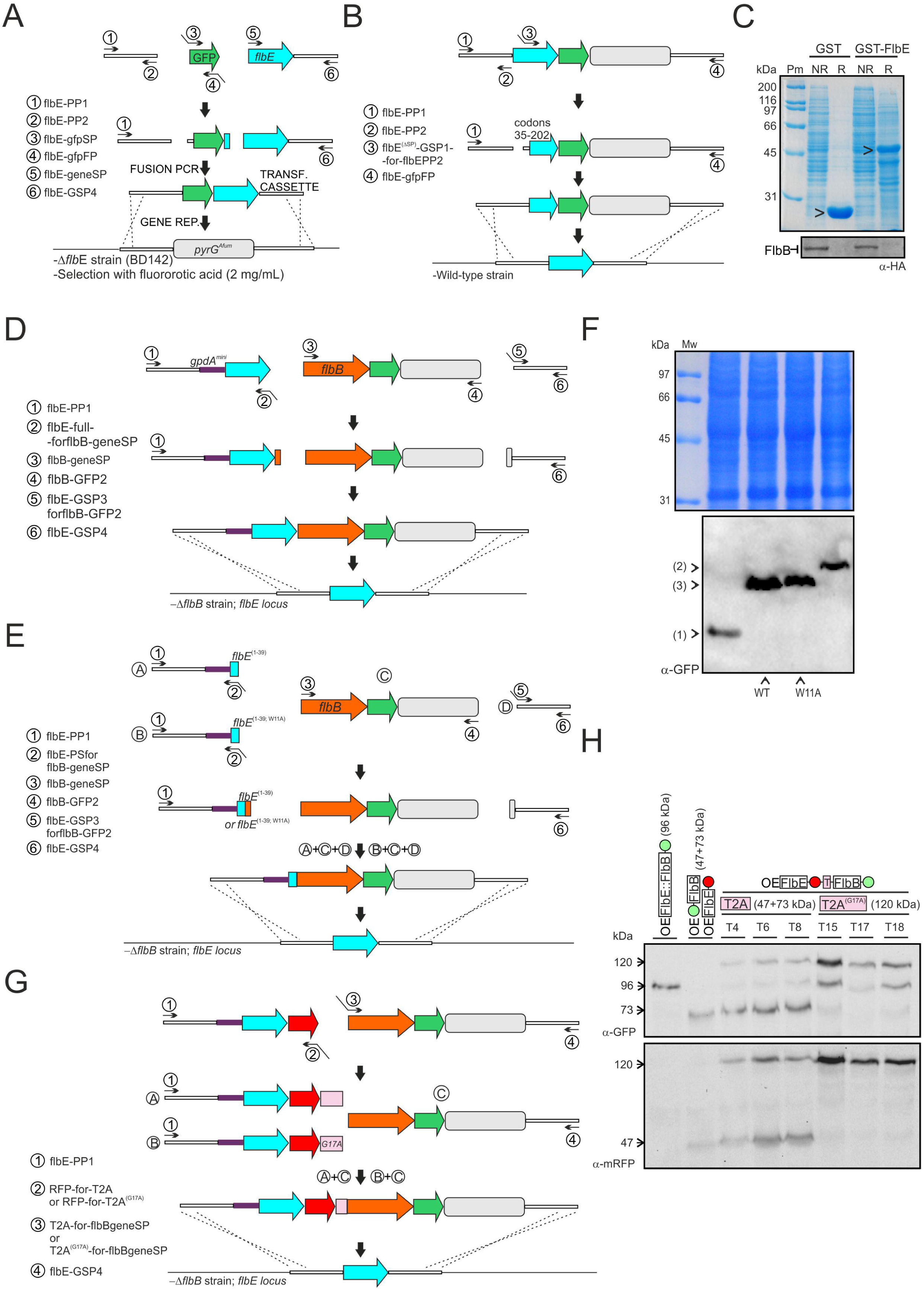
Analysis of E1 domain of FlbE. A) and B) Procedures developed for the generation of strains expressing GFP::FlbE (A) or FlbE^(34-202)^::GFP (B) chimeras, both driven by the native *flbE* promoter. Cassettes were fused and used to transform, respectively, protoplasts of a Δ*flbE* or a wild-type strain. Selection of transformants in panel A was done using fluororotic acid (2 mg/mL) [32]. C) Immunoprecipitation assay using bacterially expressed GST (negative control) and GST::FlbE forms as bait and a crude protein extract of a strain expressing an FlbB::HA_3x_ chimera. A polyacrilamide gel stained with Coomassie blue shows the concentration of each bait. D) and E) Procedure developed for the generation of strains expressing *gpdA*^*mini*^-driven FlbE::FlbB::GFP (D), as well as FlbE^(1-39)^::FlbB::GFP and FlbE^(1-39;W11A)^::FlbB::GFP chimeras (E). Cassettes for transformation were assembled by the fusion-PCR technique [62] and all of them were used to transform protoplasts of a Δ*flbB* strain. Recombination was induced at the *flbE locus*, in order to guarantee that the only source of FlbE and FlbB was the one derived from the translation of the constructs. F) Western-blot for the detection of the chimeras described in panels D and E in crude protein extracts of the corresponding strains. Polyacrilamide gels stained with Coomassie blue are shown as loading controls. G) Procedure developed for the generation of strains expressing *gpdA*^*mini*^-driven FlbE::mRFP::T2A::FlbB::GFP or FlbE::mRFP::T2A^(G17A)^::FlbB::GFP chimeras. Again, cassettes for transformation were assembled by the fusion-PCR technique [62] and all of them were used to transform protoplasts of a Δ*flbB* strain. Recombination was induced at the *flbE locus*, in order to guarantee that the only source of FlbE and FlbB was the one derived from the translation of the constructs. H) Western-blot experiment for the determination of the efficiency of the wild-type or mutant (G17A) T2A viral peptide in *Aspergillus nidulans*. Two peptides, FlbE::mRFP (47 kDa) and FlbB::GFP (73 kDa) were detected when the wild-type T2A peptide was used, together with the full length FlbE::mRFP:T2A::FlbB::GFP chimera (120 kDa). Only this latter form was detected when the mutant G17A T2A form was used. Extracts of strains expressing 1) FlbE:FlbB::GFP (96 kDa), or 2) FlbB::mRFP (47 kDa) and FlbB::GFP (73 kDa) chimeras were used as controls.

**Figure EV3.**
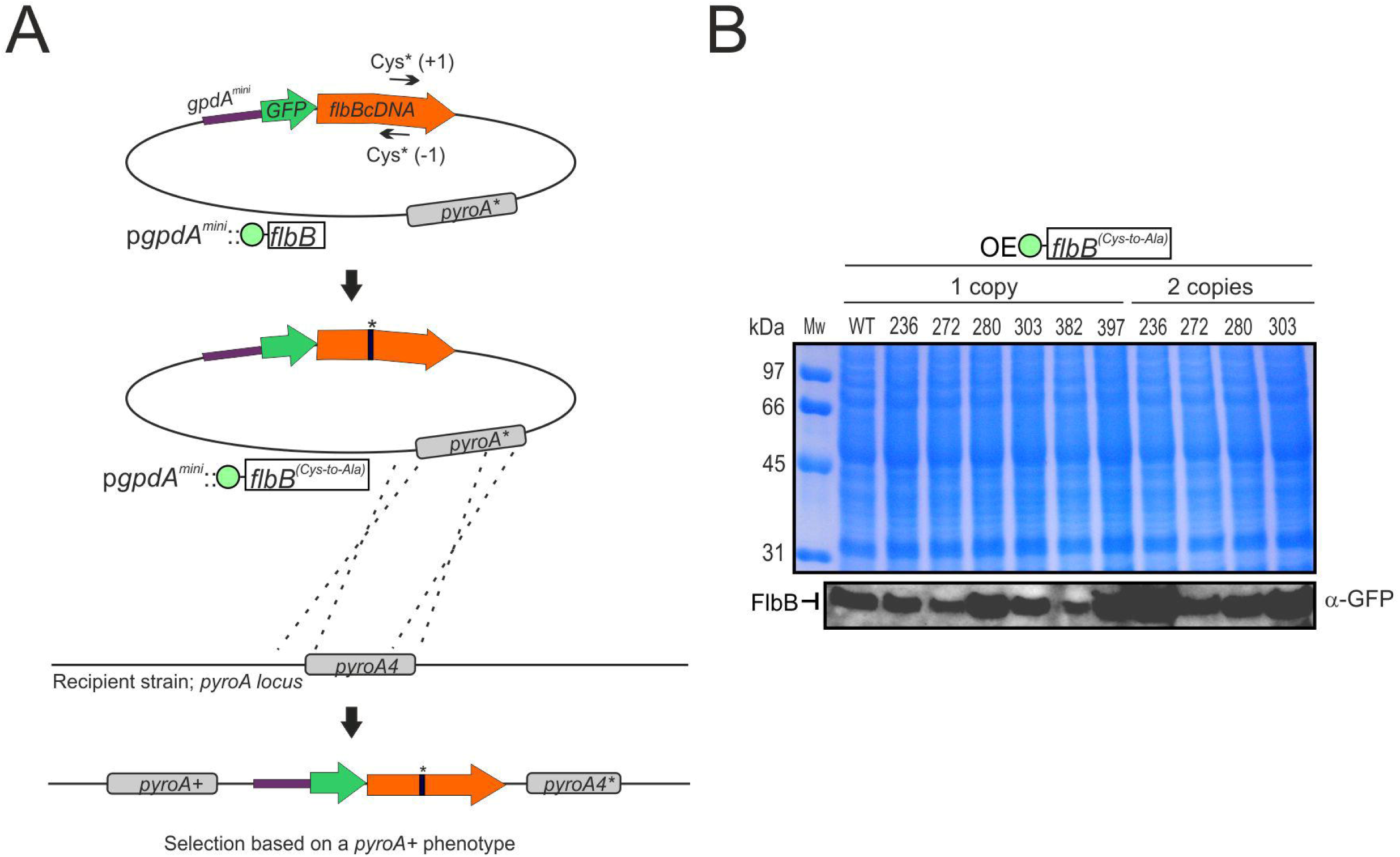
**Mutagenesis of FlbB cysteines**. A) Procedure used for the generation of *gpdA*^*mini*^-driven GFP::FlbB^(Cys-to-Ala)^ mutants [23]. A plasmid containing the fusion *gpdA*^*mini*^::*gfp*::*flbB*_*cDNA*_ was mutagenized by PCR using appropriate mutant oligonucleotides. The presence of the desired mutation was confirmed by sequencing and the mutant plasmid was used to transform protoplasts of a Δ*flbB* strain. Selection of transformants was done based on the *pyroA*^*+*^ phenotype and the correct recombination was confirmed by Southern-blot. Asterisks indicate the substitution of a codon coding for a Cys by a mutant codon coding for an Ala. B) Western-blot showing that all Cys mutants expressed GFP::FlbB^(Cys-toAla)^ chimeras of the same size. Strains expressing one or two copies of the mutant plasmid were compared. The coomassie-stained gel is shown as a loading control. See also Figure 3.

**Videos EV4: Subcellular dynamics of FlbB at hyphal tips.** Videos corresponding to three hyphae of the dual-OE strain expressing GFP::FlbB and FlbE::mRFP both driven by the *gpdA*^*mini*^ promoter. Time intervals for each frame are 0.222 (A), 0.400 (B) and 0.200 (C) seconds, respectively. Videos were generated using ImageJ (10 frames per second).

See also Figure 4.

**Figure EV5:**
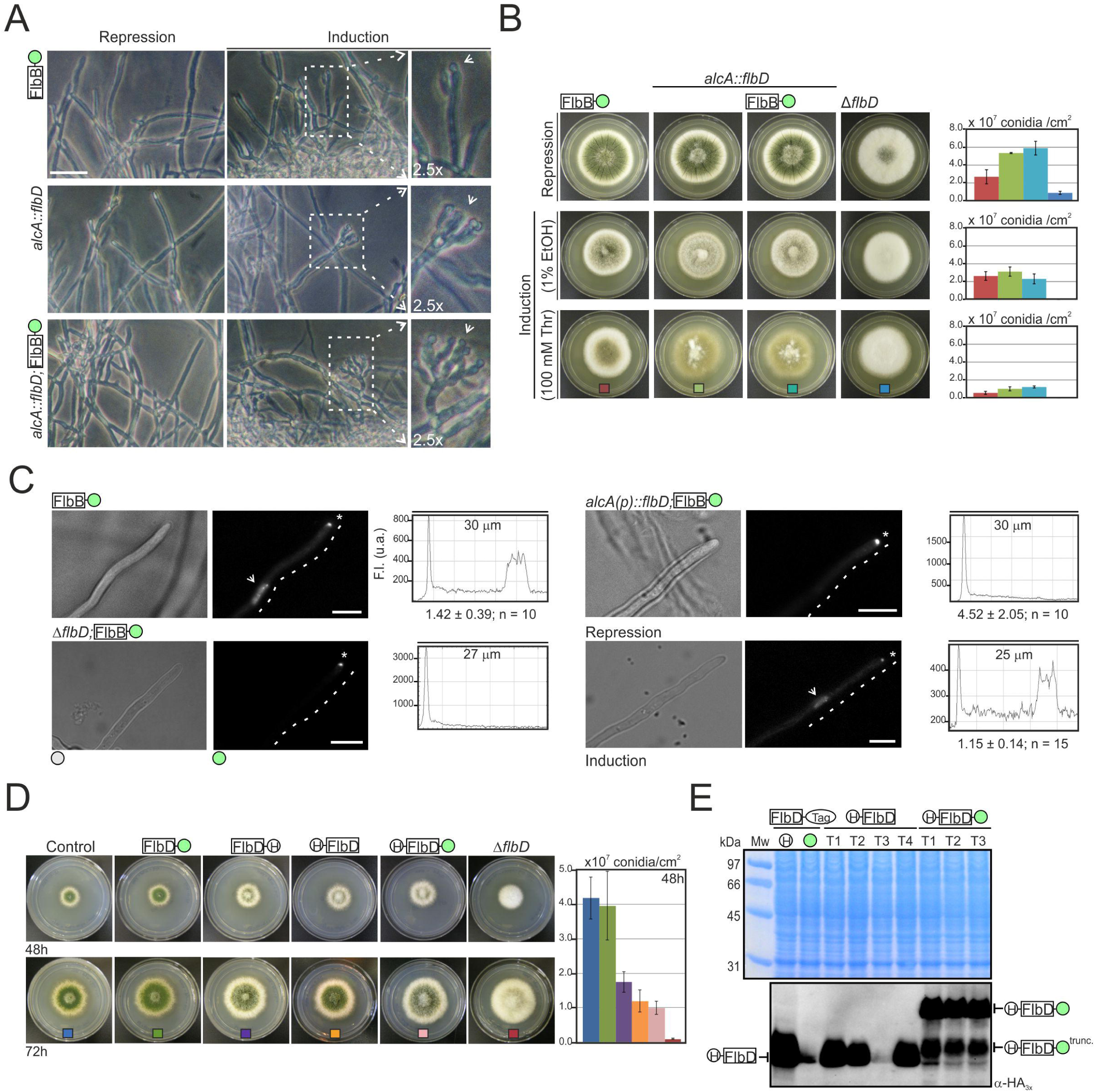
Role of FlbD in the nuclear accumulation of FlbB. A) Conidiophore development in liquid MMA as a result of *flbD* overexpression. Phenotypes of strains 1) FlbB::GFP, 2) *alcA*^*p*^::*flbD* and 3) *alcA*^*p*^::*flbD*; FlbB::GFP are shown after 18 hours of culture in MMA (2% glucose was used as the carbon source; repression of *alcA*^*p*^) (left column) or additional 22 hours of culture in MMA that contained threonine (100 mM) as the carbon source (induction of *alcA*^*p*^; middle column and 2.5x magnifications on the right). Scale bar = 25 µm. The control strain generated a limited number of conidiophores when threonine was used but those structures included only 1-3 phialides and a limited number of conidia in each phialide, resembling the morphology described by Skromne and colleagues for the simplified conidiophores produced by *A. nidulans* under carbon starvation conditions [19]. *alcA*^*p*^::*flbD* strains developed complex conidiophores. B) Effect of *flbD* overexpression on solid culture medium. 1) FlbB::GFP, 2) *alcA*^*p*^::*flbD*, 3) *alcA*^*p*^::*flbD*; FlbB::GFP and 4) Δ*flbD* strains were cultured under repressing (ACM using 2% glucose as the carbon source) (upper line) or purportedly inducing conditions (ACM plus 1% ethanol or 100 mM threonine) (middle and lower lines) for 72 hours at 37 °C. Diameter of plates is 5.5 cm. Diagrams on the right show conidia production by each strain. The values given are the mean of three replicates plus s.e.m. C) Subcellular localization of FlbB::GFP in vegetative hyphae of wild-type (up, left), Δ*flbD* (bottom, left), or *alcA*^*p*^::*flbD* (right) strains in liquid AMM. *alcA*^*p*^ was repressed by supplementing the medium with glucose (0.1 %; up) and induced by adding threonine (100 mM; bottom), respectively. The graphs show fluorescence intensity along the dotted lines. Scale bar = 5 µm. D) Phenotype of reference wild-type and null *flbD* strains, and strains expressing FlbD::GFP, FlbD::HA_3x_, HA_3x_::FlbD or HA_3x_::FlbD::GFP chimeras in AMM plates (diameter: 5.5 cm) after 48 (upper row) and 72 (lower row) hours of culture at 37 °C. The graph on the right shows conidia production by each strain after 48 hours of culture. Values given are the mean of three replicates plus s.e.m. E) Immunodetection of FlbD::HA_3x_, FlbD::GFP, HA_3x_::FlbD and HA_3x_::FlbD::GFP chimeras. The coomassie-stained gel is shown as a loading control. See also Figure 5.

