## Supplementary material for "Developmental regulators FlbE/D orchestrate the polarity site-to-nucleus dynamics of the fungal bZIP FlbB": Main Figure 5

| | |
|---|---|
| Total number of predicted TFs encoded in the *A. nidulans* genome | 490 |
| Total number of proteins predicted in the *A. nidulans* genome | 10701 |
| % of TFs | 4,6 |

| | |
|---|---|
| Number of unique proteins in *A. nidulans* with at least one LxxLL motf | 2227 |

| Keyword searches in results: | | | | | |
|---|---|---|---|---|---|
| | Number of prot with LxxLL motif | % | Num. of prot. *Anid* proteome | % | Enrichment |
| Transcription | 534 | 24,0 | 974 | 9,1 | 2,6 |
| Nucleus | 264 | 11,9 | 693 | 6,5 | 1,8 |
| Nuclear | 118 | 5,3 | 192 | 1,8 | 3,0 |
| Transcription factor | 199 | 8,9 | 402 | 3,8 | 2,4 |
| | | | | Average | 2,4 |
| Control keywords: | | | | | |
| Translation | 48 | 2,2 | 172 | 1,6 | 1,3 |
| Mitochondria | 68 | 3,1 | 378 | 3,5 | 0,9 |
| Activity | 1513 | 67,9 | 4305 | 40,2 | 1,7 |
| Metabolic | 133 | 6,0 | 479 | 4,5 | 1,3 |
| | | | | Average | 1,3 |
| | | T test | 0,007817535 | | |
