## Supplementary material for "Developmental regulators FlbE/D orchestrate the polarity site-to-nucleus dynamics of the fungal bZIP FlbB": Oligonucleotides used in this work

**Table S1. Oligonucleotides used in this study.**

| Name | Sequence (5’🡪3’) |
| --- | --- |
| flbE-PP1 | GCAACAAATTCGGCTGTTGGGCTCAGG |
| flbE-PP2’-ATG | GGTAAGGCGACGACGGCCCTCGG |
| gpdA-flbE-Up | CCGAGGGCCGTCGTCGCCTTACCCCATCCGGTGCTCTGCACTCGACC |
| gpdA-flbE-Dw | CCATCGGAAGCCGTAGAGCATGTAGACTGGCATCATGGTGATGTCTGCTCAAGCGGGG |
| flbB-geneSP | ATGACTTCGATCAGTAGTAGGCCAATACCCTTGGATCTGAAC |
| flbE-GSP2 | CGAAAACGTTTTGTTGAAGAATCGCGTTAATGTCTC |
| flbE-GFP1 | GAGACATTAACGCGATTCTTCAACAAAACGTTTTCGGGAGCTGGTGCAGGCGCTGGAGCC |
| FlbE-GFP2 | TCATCAGTCGTAATATAACTCGTACAGCAATCAGTCTGAGAGGAGGCACTGATGCG |
| flbE-GSP3 | TGATTGCTGTACGAGTTATATTACGACTGATGA |
| flbE-GSP4 | GCTTACCTGCTGGATCTCCTGCCGGTACTTAGG |
| FlbEW11AUp | CTCTACGGCTTCCGAGCGCCCCGAGCTGG |
| FlbEW11ADw | CCAGCTCGGGGCGCTCGGAAGCCGTAGAG |
| FlbEK51AUp | GGACTCATTTAGGGCGACGGAGCCG |
| FlbEK51ADw | CGGCTCCGTCGCCCTAAATGAGTCC |
| FlbED70A;D73AUp | CGCTTCATTGAACAGTATGCCCCCGAGGCCGAAAGC |
| FlbED70A;D73ADw | GCTTTCGGCCTCGGGGGCATACTGTTCAATGAAGCG |
| FlbEY85A;V86AUp | GCCTTATGCTGCTGCTGCTGCGAAAACG |
| FlbEY85A;V86ADw | CGTTTTCGCAGCAGCAGCAGCATAAGGC |
| FlbEP182AUp | GGGTCGTCTAGTAACACAGCGTCGACGCCG |
| FlbEP182Dw | CGGCGTCGACGCTGTGTTACTAGACGACCC |
| 0721-PP2 | CATGGTAAGGCGACGACGGCCCTC |
| 0721-gfpSP | GAGGGCCGTCGTCGCCTTACCATGAGTAAAGGAGAAGAACTTTTCACTGGAGTT |
| 0721-gfpFP | CCATCGGAAGCCGTAGAGCATGTAGACTGGCATTTTGTATAGTTCATCCATGCCATGTGT |
| FlbE-GSP1(ΔSP)forFlbEPP2 | CGAGGGCCGTCGTCGCCTTACCATGGAATACATACAAAGACCGATAACGAACAAGTCG |
| FlbE-SPforFlbBgeneSP | GTTCAGATCCAAGGGTATTGGCCTACTACTGATCGAAGTCATCGGTCTTTGTATGTATTCCGCCGTAGC |
| FlbEfullforFlbBgeneSP | GTTCAGATCCAAGGGTATTGGCCTACTACTGATCGAAGTCATCGAAAACGTTTTGTTGAAGAATCGCGTTAATGTCTC |
| FlbE-GSP3forFlbBGFP2 | TGACCTGACAGCTCGCTTTTTTTCTGAGCTTTCTAATGCTGATTGCTGTACGAGTTATATTACGACTGATGA |
| flbB-geneSP | ATGACTTCGATCAGTAGTAGGCCAATACCCTTGGATCTGAAC |
| flbB-GFP2 | GCATTAGAAAGCTCAGAAAAAAAGCGAGCTGTCAGGTCAGTCTGAGAGGAGGCACTGATGCG |
| RFPDw-for-T2A | AGGTCCAGGATTCTCCTCGACGTCACCGCATGTTAGCAGACTTCCTCTGCCCTCGGCGCCGGTGGAGTGGCG |
| RFPDw-for-T2A* | AGGTGCAGGATTCTCCTCGACGTCACCGCATGTTAGCAGACTTCCTCTGCCCTCGGCGCCGGTGGAGTGGCG |
| T2A-for-flbBgeneSP | GAGGGCAGAGGAAGTCTGCTAACATGCGGTGACGTCGAGGAGAATCCTGGACCTATGACTTCGATCAGTAGTAGGCCAATACCCTTGGATCTGAAC |
| T2A*-for-flbBgeneSP | GAGGGCAGAGGAAGTCTGCTAACATGCGGTGACGTCGAGGAGAATCCTGCACCTATGACTTCGATCAGTAGTAGGCCAATACCCTTGGATCTGAAC |
| myoE-PP1 | GTTGCGACGGATATCGAGGAACG |
| myoE-PP2 | CATCGTGAACACGAAAACCGCTGC |
| myoE-GSP3 | TAATATCTCTCAGTTTGTCCACTACCACCGCTGG |
| myoE-GSP4 | CTGCTGCAGGGGATACCATGAAACG |
| myoE-SMP1 | GCAGCGGTTTTCGTGTTCACGATGACCGGTCGCCTCAAACAATGCTCT |
| myoE-GFP2 | CCAGCGGTGGTAGTGGACAAACTGAGAGATATTAGTCTGAGAGGAGGCACTGATGCG |
| flbB-C236A up | GCTACAGATCCGGCTGATGCACTTGCCG |
| flbB-C236A dw | CGGCAAGTGCATCAGCCGGTACTGTAGC |
| flbB-C272A up | CTGGAATCACCCGCTCGAGATCATACCG |
| flbB-C272A dw | CGGTATGATCTCGAGCGGGTGATTCCAG |
| flbB-C280A up | CCGACTACCTCGCCCGACGATCCATC |
| flbB-C280A dw | GATGGATCGTCGGGCGAGGTAGTCGG |
| flbB-C303A up | GCTGATGGCCACCGCCCCACCGCCAAGC |
| flbB-C303A dw | GCTTGGCGGTGGGGCGGTGGCCATCAGC |
| flbB_C382A(+1) | GAGAACAAGGTGCGCGCATACGGATTCGGG |
| flbB_C382A(-1) | CCCGAATCCGTATGCGCGCACCTTGTTCTC |
| FlbB(sek3) | ggcgctggaggacaggc |
| FlbB(sek5) | gcgttgcgttttcccggc |
| flbB-NLS-1 | GGGTTTTGGTCCTCTTGCTGCGGCAGGTTGCCC |
| flbB-PP1 | GTTTTCTGGTCCTCGGTCAACCGGTGG |
| flbB-GSP4 | GAAAGGTGCGTGGGTTCGAATCCCACC |
| FlbD-PP1 | CGACTCCATGACCACATTCC |
| FlbD-PP2 | CATTTGCGAAACTGTGTTGG |
| FlbD-geneSP | ATGGCTCCAACACACCGTCG |
| FlbD-GSP4 | GGCAGGCGTTACTGTATACGT |
| FlbD-HA-SP | CCAACACAGTTTCGCAAATGGGCCGCATCTTTTACCCATACG |
| FlbD-HA-FP | CGACGGTGTGTTGGAGCCATCTGAGCAGCGTAATCTGGAACG |
| hhoA-GSP5 | CTACAGTTGAGCCTATGCACAAG |
| hhoA-GSP6 | GGCCTTCTTGTTCTTAGCAGTC |
| hhoA-GSP3 | ACAGGCAGCCTGGCTTGG |
| hhoA-GSP4 | GCAACAGTCGACAGCACAGC |
| hhoA-GSP3´ | CCAAGCCAGGCTGCCTGTCTGTCTGAGAGGAGGCACTGATGC |
| hhoA-GSP6´ | GACTGCTAAGAACAAGAAGGCCGGAGCTGGTGCAGGCGCTGGAGCC |
| FlbD-LigBox*-Up | CGGATGGGCGCTGACAACGCTTTGAAC |
| FlbD-LigBox*-Dw | GTTCAAAGCGTTGTCAGCGCCCATCCG |
| FlbDsec | TCACGGTCTACTTGCACC |
| GFP(NFlbD)sec | CACTAGTTGGAGAAGCAGC |
| FlbD-Cterm | CGACAACCTCTTGAACTGAACGATCACACGACTC |
| cMybSTOP | GAGTCGTGTGATCGTTCAGAGACCTCTCTTCTTGC |
| FlbB_L333AL334A_Up | CCTCACAACGGCAGCCAATCTTAGC |
| FlbB_L333AL334A_Dw | GCTAAGATTGGCTGCCGTTGTGAGG |
