## Supplementary material for "Developmental regulators FlbE/D orchestrate the polarity site-to-nucleus dynamics of the fungal bZIP FlbB": Strains of A. nidulans used in this work

**Table S1: *A. nidulans* strains used in this study.**

| Strain | Genotype | Source |
| --- | --- | --- |
| BD142 | *pyrG89; argB2; pyroA4,* Δ*nkuA::argB;* Δ*flbE::pyrG^Af^, veA1* | (Garzia *et al.*, 2009) |
| BD167 | *pyrG89; argB2; flbB::gfp::pyrG^Af^, pyroA4,* ∆*nkuA::argB; veA1* | (Etxebeste *et al.*, 2008) |
| BD177 | *pyrG89; argB2;* ∆*flbB::riboB^Af^, pabaB22, pyroA4,* ∆*nkuA::argB; riboB2, veA1* | (Garzia *et al.*, 2009) |
| BD178 | *pyrG89; argB2; pabaB22, pyroA4,* Δ*nkuA::argB; riboB2,* Δ*flbE::riboB^Af^, veA1.* | (Garzia *et al.*, 2009) |
| BD185 | *pyrG89; argB2; gfp::flbB, pyroA4,* Δ*nkuA::argB; veA1* | (Etxebeste *et al.*, 2008) |
| BD186 | *pyrG89; argB2; pyroA4, ΔnkuA::argB; flbE::gfp::pyrG^Af^, veA1* | (Garzia *et al.*, 2009) |
| BD197 | *pyrG89; argB2; pyroA4,* Δ*nkuA::argB; veA1,* Δ*flbD::pyrG^Af^* | (Garzia *et al.*, 2010) |
| BD198 | *pyrG8; argB2; pabaB22, pyroA4,* ∆*nkuA::argB; veA1, riboB2,* ∆*flbD::riboB^Af^* | (Garzia *et al.*, 2010) |
| BD207 | *pyrG89; argB2; gfp::flbB, pyroA4,* Δ*nkuA::argB; flbE::mCh::pyroA, veA1* | (Garzia *et al.*, 2009) |
| BD209 | *pyrG89; argB2; flbB::gfp::pyrG^Af^, pabaB22, pyroA4,* Δ*nkuA::argB; veA1, riboB2,* Δ*flbD::riboB^Af^* | (Garzia *et al.*, 2010) |
| BD246 | *pyrG89; argB2; pyroA4,*  Δ*nkuA::argB; veA1, flbD::gfp::pyrG^Af^* | (Garzia *et al.*, 2010) |
| BD271 | *pyrG89; argB2; pabaB22, pyroA^+^::[alcA(p)::flbD]::pyroA4*,* Δ*nkuA::argB; veA1, riboB2,* Δ*flbD::riboB^Af^* | (Garzia *et al.*, 2010) |
| BD294 | *pyrG89; argB2; flbB::HA_3x_::pyrG^Af^, pyroA4, ΔnkuA::argB; veA1* | (Herrero-Garcia *et al.*, 2015) |
| BD375 | *pyrG89; argB2; pyroA4,* ∆*nkuA::argB; veA1, flbD::HA_3x_::pyrG^Af^* | (Garzia *et al.*, 2010) |
| BD630 | *pyrG89; argB2;* Δ*flbB::riboB^Af^, pabaB22, pyroA^+^::[gpdA^mini^::gfp::flbB]_1x_::pyroA4*,* Δ*nkuA::argB; riboB2, veA1* | (Perez-de-Nanclares-Arregi and Etxebeste, 2014) |
| BD643 | *pyrG89; argB2;* Δ*flbB::riboB^Af^, pabaB22, pyroA^+^::[gpdA^mini^::gfp::flbB^(C397A)^]_1x_::pyroA4*,* Δ*nkuA::argB; riboB2, veA1* | (Herrero-Garcia *et al.*, 2015) |
| BD721 | *pyrG89; argB2;* Δ*flbB::riboB^Af^, pabaB22, pyroA^+^::[gpdA^mini^::gfp::flbB]_1x_::pyroA4*,* Δ*nkuA::argB; gpdA^mini^::flbE::mrfp::pyrG^Af^, veA1* | (Herrero-Garcia *et al.*, 2015) |
| BD723 | *pyrG89; argB2;* Δ*flbB::riboB^Af^, pabaB22, pyroA^+^::[gpdA^mini^::gfp::flbB^(C382A)^]_1x_::pyroA4*,* Δ*nkuA::argB; gpdA^mini^::flbE::mrfp::pyrG^Af^, veA1* | (Herrero-Garcia *et al.*, 2015) |
| BD731 | *pyrG89; argB2; pyroA4, ΔnkuA::argB; flbE^(34-202)^::gfp::pyrG^Af^, veA1* | This study |
| BD735 | *pyrG89; argB2; pyroA4,* Δ*nkuA::argB; gfp::flbE, veA1* | This study |
| BD770 | *pyrG89; argB2; pyroA4,* Δ*nkuA::argB; gpdA^mini^::flbE::mrfp::pyrG^Af^, veA1* | This study |
| BD773 | *pyrG89; argB2; pyroA4;* Δ*nkuA::argB; gpdA^mini^::flbE::gfp::pyrG^Af^, veA1* | This study |
| BD841 | *pyrG89; argB2;* Δ*flbB::riboB^Af^, pabaB22, pyroA^+^::[gpdA^mini^::gfp::flbB]_1x_::pyroA4*,* Δ*nkuA::argB; riboB2, gpdA^mini^::flbE::stag::pyrG^Af^, veA1* | This study |
| BD845 | *pyrG8; argB2; flbB::gfp::pyrG^Af^, pabaB22, pyroA4,* Δ*nkuA::argB; hhoA::mCh::pyroA; veA1, riboB2,* Δ*flbD::riboB^Af^* | This study |
| BD852 | *pyrG89; argB2; flbB::gfp::pyrG^Af^, pabaB22, pyroA^+^::[alcA(p)::flbD]::pyroA4*,* Δ*nkuA::argB; veA1, riboB2,* Δ*flbD::riboB^Af^* | This study |
| BD918 | *pyrG89; argB2;* Δ*flbB::riboB^Af^, pabaB22, pyroA^+^::[gpdA^mini^::gfp::flbB^(C272A)^]_1x_::pyroA4*,* Δ*nkuA::argB; riboB2, veA1* | This study |
| BD920 | *pyrG89; argB2;* Δ*flbB::riboB^Af^, pabaB22, pyroA^+^::[gpdA^mini^::gfp::flbB^(C272A)^]_2x_::pyroA4*,* Δ*nkuA::argB; riboB2, veA1* | This study |
| BD921 | *pyrG89; argB2;* Δ*flbB::riboB^Af^, pabaB22, pyroA^+^::[gpdA^mini^::gfp::flbB^(C280A)^]_1x_::pyroA4*,* Δ*nkuA::argB; riboB2, veA1* | This study |
| BD923 | *pyrG89; argB2;* Δ*flbB::riboB^Af^, pabaB22, pyroA^+^::[gpdA^mini^::gfp::flbB^(C280A)^]_2x_::pyroA4*,* Δ*nkuA::argB; riboB2, veA1* | This study |
| BD925 | *pyrG89; argB2;* Δ*flbB::riboB^Af^, pabaB22, pyroA^+^::[gpdA^mini^::gfp::flbB^(C236A)^]_1x_::pyroA4*, ΔnkuA::argB; riboB2, veA1* | This study |
| BD930 | *pyrG89; argB2;* Δ*flbB::riboB^Af^, pabaB22, pyroA^+^::[gpdA^mini^::gfp::flbB^(C236A)^]_2x_::pyroA4*,* Δ*nkuA::argB; riboB2, veA1* | This study |
| BD936 | *pyrG89; argB2;* Δ*flbB::riboB^Af^, pabaB22, pyroA^+^::[gpdA^mini^::gfp::flbB^(C303A)^]_1x_::pyroA4*,* Δ*nkuA::argB; riboB2, veA1* | This study |
| BD941 | *pyrG89; argB2;* Δ*flbB::riboB^Af^, pabaB22, pyroA^+^::[gpdA^mini^::gfp::flbB^(C303A)^]_2x_::pyroA4*,* Δ*nkuA::argB; riboB2, veA1* | This study |
| BD995 | *pyrG89; argB2; pyroA4,* Δ*nkuA::argB; gpdA^mini^::flbE^(W11A)^::gfp::pyrG^Af^, veA1* | This study |
| BD997 | *pyrG89; argB2; pyroA4,* Δ*nkuA::argB; gpdA^mini^::flbE^(K51A)^::gfp::pyrG^Af^, veA1* | This study |
| BD998 | *pyrG89; argB2; pyroA4,* Δ*nkuA::argB; gpdA^mini^::flbE^(Y85A;V86A)^::gfp::pyrG^Af^, veA1* | This study |
| BD1000 | *pyrG89; argB2; pyroA4,* Δ*nkuA::argB; gpdA^mini^::flbE^(P182A)^::gfp::pyrG^Af^, veA1* | This study |
| BD1001 | *pyrG89; argB2;* Δ*flbB::riboB^Af^, pabaB22, pyroA^+^::[gpdA^mini^::gfp::flbB]_1x_::pyroA4*,* Δ*nkuA::argB; riboB2, gpdA^mini^::flbE^(W11A)^::mrfp::pyrG^Af^, veA1* | This study |
| BD1005 | *pyrG89; argB2;* Δ*flbB::riboB^Af^, pabaB22, pyroA^+^::[gpdA^mini^::gfp::flbB]_1x_::pyroA4*,* Δ*nkuA::argB; riboB2, gpdA^mini^::flbE^(K51A)^::mrfp::pyrG^Af^, veA1* | This study |
| BD1012 | *pyrG89; argB2;* Δ*flbB::riboB^Af^, pabaB22, pyroA^+^::[gpdA^mini^::gfp::flbB]_1x_::pyroA4*,* Δ*nkuA::argB; riboB2, gpdA^mini^::flbE^(D70A;D73A)^::mrfp::pyrG^Af^, veA1* | This study |
| BD1014 | *pyrG89; argB2;* Δ*flbB::riboB^Af^, pabaB22, pyroA^+^::[gpdA^mini^::gfp::flbB]_1x_::pyroA4*,* Δ*nkuA::argB; riboB2, gpdA^mini^::flbE^(Y85A;V86A)^::mrfp::pyrG^Af^, veA1* | This study |
| BD1017 | *pyrG89; argB2;* Δ*flbB::riboB^Af^, pabaB22, pyroA^+^::[gpdA^mini^::gfp::flbB]_1x_::pyroA4*,* Δ*nkuA::argB; riboB2, gpdA^mini^::flbE^(P182A)^::mrfp::pyrG^Af^, veA1* | This study |
| BD1020 | *pyrG89; argB2;* Δ*flbB::riboB^Af^, pabaB22, pyroA4,* Δ*nkuA::argB; gpdA^mini^::flbE^(1-39)^::flbB^(1-426)^::gfp::pyrG^Af^, veA1* | This study |
| BD1023 | *pyrG89; argB2;* Δ*flbB::riboB^Af^, pabaB22, pyroA4,* Δ*nkuA::argB; gpdA^mini^::flbE^(1-39,W11A)^::flbB^(1-426)^::gfp::pyrG^Af^, veA1* | This study |
| BD1026 | *pyrG89; argB2;* Δ*flbB::riboB^Af^, pabaB22, pyroA4,* Δ*nkuA::argB; gpdA^mini^::flbE^(1-202)^::flbB^(1-426)^::gfp::pyrG^Af^, veA1* | This study |
| BD1029 | *pyrG89; argB2;* Δ*flbB::riboB^Af^, pabaB22, pyroA^+^::[gpdA^mini^::gfp::flbB^(C236A)^]_1x_::pyroA4*,* Δ*nkuA::argB; riboB2, gpdA^mini^::flbE::mrfp::pyrG^Af^, veA1* | This study |
| BD1037 | *pyrG89; argB2;* Δ*flbB::riboB^Af^, pabaB22, pyroA^+^::[gpdA^mini^::gfp::flbB^(C272A)^]_1x_::pyroA4*,* Δ*nkuA::argB; riboB2, gpdA^mini^::flbE::mrfp::pyrG^Af^, veA1* | This study |
| BD1047 | *pyrG89; argB2;* Δ*flbB::riboB^Af^, pabaB22, pyroA^+^::[gpdA^mini^::gfp::flbB^(C272A;C382A)^]_1x_::pyroA4*,* Δ*nkuA::argB; riboB2, gpdA^mini^::flbE::mrfp::pyrG^Af^, veA1* | This study |
| BD1057 | *pyrG89; argB2;* Δ*flbB::riboB^Af^, pabaB22, pyroA^+^::[gpdA^mini^::gfp::flbB^(C272A;C382A)^]_1x_::pyroA4*,* Δ*nkuA::argB; riboB2, veA1* | This study |
| BD1058 | *pyrG89; argB2;* Δ*flbB::riboB^Af^, pabaB22, pyroA^+^::[gpdA^mini^::gfp::flbB^(C272A;C382A)^]_2x_::pyroA4*,* Δ*nkuA::argB; riboB2, veA1* | This study |
| BD1060 | *pyrG89; argB2; pyroA4,* Δ*nkuA::argB, gpdA^mini^::flbE^(D70A;D73A)^::gfp::pyrG^Af^, veA1* | This study |
| BD1093 | *pyrG89; argB2; pyroA4,* Δ*nkuA::argB; veA1, HA_3x_::flbD* | This study |
| BD1096 | *pyrG89; argB2; pyroA4,* Δ*nkuA::argB; veA1, HA_3x_::flbD::gfp::pyrG^Af^* | This study |
| BD1111 | *pyrG89; argB2; gfp::flbB, pyroA4,* Δ*nkuA::argB; veA1, gpdA^mini^::flbE^(D70A;D73A)^::mrfp::pyrG^Af^* | This study |
| BD1137 | *pyrG89; argB2;* Δ*flbB::riboB^Af^, pabaB22, pyroA4,* Δ*nkuA::argB; gpdA^mini^::flbE^(1-202)^::mrfp::t2a::flbB^(1-426)^::gfp::pyrG^Af^, veA1* | This study |
| BD1140 | *pyrG89; argB2;* Δ*flbB::riboB^Af^, pabaB22, pyroA4,* Δ*nkuA::argB; gpdA^mini^::flbE^(1-202)^::mrfp::t2a^(G17A)^:flbB^(1-426)^::gfp::pyrG^Af^, veA1* | This study |
| BD1148 | *pyrG8; argB2; pabaB22, pyroA4,* ∆*nkuA::argB; veA1, riboB2, flbD^(L309A;L312A)^::gfp::pyrG^Af^* | This study |
| BD1150 | *pyrG89; argB2; pyroA4,* Δ*nkuA::argB; veA1, flbD^(L309A;L312A)^* | This study |
| BD1168 | *pyrG89; argB2; flbB::gfp::pyrG^Af^, pyroA4,* Δ*nkuA::argB; veA1, flbD^(L309A;L312A)^* | This study |
| BD1186 | *pyrG89; argB2; flbB^(L333A;L334A)^, pyroA4,* Δ*nkuA::argB; veA1* | This study |
| BD1188 | *pyrG89; argB2; gfp::flbB^(L333A;L334A)^, pyroA4,* Δ*nkuA::argB; veA1* | This study |
| BD1190 | *pyrG89; argB2; pyroA4,* Δ*nkuA::argB; veA1, flbD^(1-112)^* | This study |
| BD1216 | *pyrG8; argB2; pabaB22, pyroA4,* ∆*nkuA::argB; veA1, riboB2, flbD^(E14G;R87Q)^* | This study |
| BD1218 | *pyrG8; argB2; pabaB22, pyroA4,* ∆*nkuA::argB; veA1, riboB2, flbD5´-UTR(^A-729G)^::flbD::gfp::pyrG^Af^* | This study |
| BD1226 | *pyrG89; argB2; pyroA4,* Δ*nkuA::argB; veA1, flbD5´-UTR(^A-729G)^::flbD* | This study |
| BD1342 | *pyrG89; argB2; flbB::gfp::pyrG^Af^, pyroA4,* Δ*nkuA::argB; veA1, flbD^(E14G;R87Q)^* | This study |
| BD1340 | *pyrG89; argB2; flbB::gfp::pyrG^Af^, pyroA4,* Δ*nkuA::argB; veA1, flbD^(1-112)^* | This study |
| BD___ | *pyrG89; argB2; flbB::gfp::pyrG^Af^, pyroA4,* Δ*nkuA::argB; veA1, flbD5´-UTR(^A-729G)^::flbD* | This study |
| MAD2698 | *pyrG89, pabaA1; GFP::FlbB; chaA1, nudA1, veA1* | This study |
| MAD2949 | *pyrG89; argB2,* Δ*myoE::pyrG^Af^*; *gfp::flbB, pyroA4,* Δ*nkuA::argB;, veA1* | This study |
| MAD4328 | *pyrG89; argB2;* Δ*flbB::riboB^Af^, pabaB22, pyroA^+^::[gpdA^mini^::gfp::flbB^(C382A)^]_1x_::pyroA4*,* Δ*nkuA::argB; riboB2, veA1* | (Herrero-Garcia *et al.*, 2015) |
| TN02A3 | *pyrG89; argB2; pyroA4,* Δ*nkuA::argB; veA1* | (Nayak *et al.*, 2006) |
