## Supplementary figures and images for "Developmental regulators FlbE/D orchestrate the polarity site-to-nucleus dynamics of the fungal bZIP FlbB"

### Main Figure 5

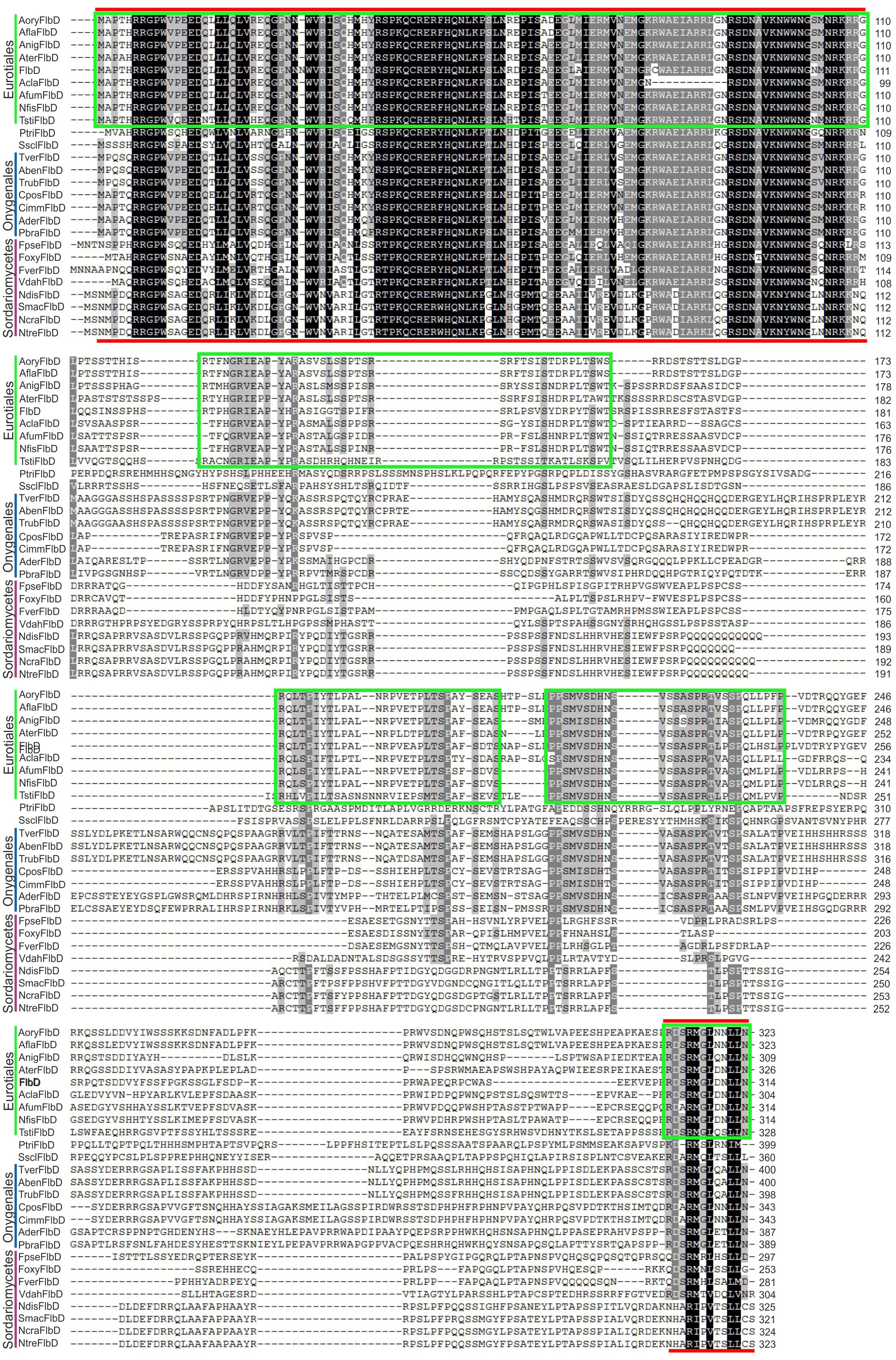
